# Live imaging reveals oocyte membrane pre-remodeling forming permissive space for asymmetric division

**DOI:** 10.64898/2026.10.14.759078

**Authors:** Kaiying Geng, Xindi Hu, Peng Fu, Yingnan Bo, Ge Wang, Xuebing Yang, Jing Liang, Kaixin Cheng, Nan Sun, Zixiong Wang, Chen Yang, Xiaoyun Zhao, Zhen Li, Hua Zhang, Yan Zhang

## Abstract

Successful oocyte maturation requires first polar body (PB1) extrusion, yet how the oocyte prepares for this asymmetric event remains unclear. Live membrane imaging of mouse oocytes reveals pronounced regional heterogeneity in plasma-membrane dynamics, generating an early polarized pattern across the surface. Whereas most of the oocyte surface and its microvilli remain relatively stable, a discrete membrane domain collapses and retracts around germinal vesicle breakdown (GVBD), separating from the zona pellucida to establish a pre-extrusion space (PES). Transient oocyte comb-like protrusions arise within the PES and subsequently collapse, converting the protrusion-filled region into an open cavity before PB1 extrusion. Mechanistically, early ERK-dependent dissociation of the cortical scaffold ZO-1 is required for this space to form. ERK2 phosphorylates ZO-1, suppresses its phase separation, and promotes its removal from the cortex. Blocking MEK-ERK signaling prevents ZO-1 dissociation, abolishes pre-extrusion space formation, and impairs PB1 extrusion, whereas ERK activation restores these events in aged oocytes. Together, our findings identify a membrane collapse-based pre-remodeling mechanism that licenses PB1 extrusion and reveal ERK-ZO-1 signaling as a key regulator of oocyte maturation.

## Introduction

Asymmetric division during mammalian oocyte maturation is a remarkable morphogenetic process. It preserves the bulk of the maternal cytoplasm within the female gamete while segregating one set of homologous chromosomes into a diminutive PB1^1–4^. This asymmetry is essential for oocyte quality and subsequent embryogenesis. It is achieved through coordinated spindle organization and migration^5–8^, cortical polarization^9^ and actin-dependent force generation^10–12^. Despite substantial progress in understanding these late meiotic events, how the oocyte spatially prepares its surface before PB1 extrusion remains unknown.

Mammalian oocytes face a distinctive spatial challenge during asymmetric division. Unlike most dividing cells, the oocyte is enclosed by the zona pellucida (ZP), a mechanically confining extracellular glycoprotein matrix that remains closely apposed to the oocyte surface^13–15^. During follicular growth, the oocyte-ZP interface is densely occupied by oocyte microvilli (Oo-Mvi) and granulosa cell-derived transzonal projections (GC-TZPs)^16–18^. Together with membrane-associated junctional scaffolds, these structures maintain structural and metabolic communication between the oocyte and its surrounding somatic cells^19–22^. This compact interface leaves limited physical clearance for membrane deformation and PB1 extrusion. The oocyte must therefore locally reorganize its membrane and disengage from the ZP to create sufficient clearance for PB1 extrusion. When and how this membrane remodeling is established remain unknown.

Previous studies have established that late meiosis I (MI) is accompanied by pronounced cortical actin remodeling^23–25^. This remodeling contributes to spindle positioning^11, 12^ and helps shape the site of PB1 extrusion^26^. This late cortical domain coordinates spindle positioning, membrane protrusion and cytokinetic asymmetry^27, 28^. However, these events occur several hours after GVBD and primarily describe the execution phase of PB1 extrusion rather than the initial preparation of the oocyte surface. Whether the oocyte undergoes an earlier cytological remodeling event to disengage its membrane from the surrounding ZP and establish a permissive space for PB1 extrusion remains unknown.

The generation of a locally permissive membrane domain is likely to require coordinated remodeling of membrane-cortex interactions and membrane-associated scaffolds^29^. Such remodeling may involve dynamic reorganization of proteins that couple the plasma membrane to the underlying cortical cytoskeleton^24, 30^. Although best known as a tight-junction protein, zonula occludens-1 (ZO-1) is a multidomain scaffold that can link cadherin-based adhesion complexes to cortical actin even in cells lacking tight junctions^31–33^. Consistent with this broader scaffolding role, ZO-1 localizes to the oocyte-granulosa cell interface despite the absence of functional tight junctions in mouse ovarian follicles^34^. ZO-1 can undergo phase separation, and this process is sensitive to phosphorylation, suggesting a potential mechanism for the rapid reorganization of ZO-1 assemblies^35, 36^. Extracellular signal-regulated kinase (ERK) is strongly activated following meiotic resumption and regulates multiple aspects of oocyte maturation^37^. Whether ERK-dependent remodeling of ZO-1 contributes to the early spatial organization of the oocyte surface, however, remains unknown.

Here, we identify an early membrane-remodeling program that spatially prepatterns the site of PB1 extrusion. High-resolution live imaging of oocytes expressing a membrane-targeted fluorescent reporter revealed that around GVBD, the plasma membrane undergoes localized retraction from the ZP to generate a pre-extrusion space (PES), well before formation of the late meiotic cortical actin domain. PES formation is accompanied by the transient appearance of parallel, comb-like membrane protrusions that project across the widening oocyte-ZP space and subsequently collapse, leaving an open cavity. The PES marks the predominant site of PB1 extrusion, and its area is strongly correlated with the area of the subsequently extruded PB1. We further show that ERK2 phosphorylation suppresses ZO-1 phase separation in vitro and that ERK-dependent ZO-1 remodeling supports localized membrane retraction and PES formation in oocytes. This pathway is compromised in oocytes from aged females, in which defective ZO-1 dissociation is associated with diminished PES formation and reduced PB1 extrusion. Pharmacological activation of ERK restores ZO-1 remodeling and PES formation and increases PB1 extrusion in aged oocytes. These findings identify a previously unrecognized membrane pre-remodeling event that precedes canonical cortical polarization and establishes a spatially permissive domain for asymmetric oocyte division.

## Results

### High-resolution live imaging reveals local membrane remodeling that establishes a pre-extrusion space during early oocyte maturation

To characterize plasma membrane dynamics during oocyte maturation, we performed high-resolution live-cell imaging using oocytes from *Oct4-CreER^T2^;mTmG* mice^38, 39^, in which the oocyte plasma membrane was genetically labelled with GFP (Fig. S1A). Following tamoxifen induction, Cre-mediated recombination switched the membrane-targeted fluorescent reporter from tdTomato to GFP, enabling specific visualization of oocyte plasma membrane and membrane-derived subcellular structures (Fig. S1B-C). We then performed high-resolution three-dimensional time-lapse imaging over 16 h of in vitro maturation at 2-h intervals to monitor membrane dynamics throughout meiotic progression (Fig. 1A and Movie S1).

**Fig. 1.**
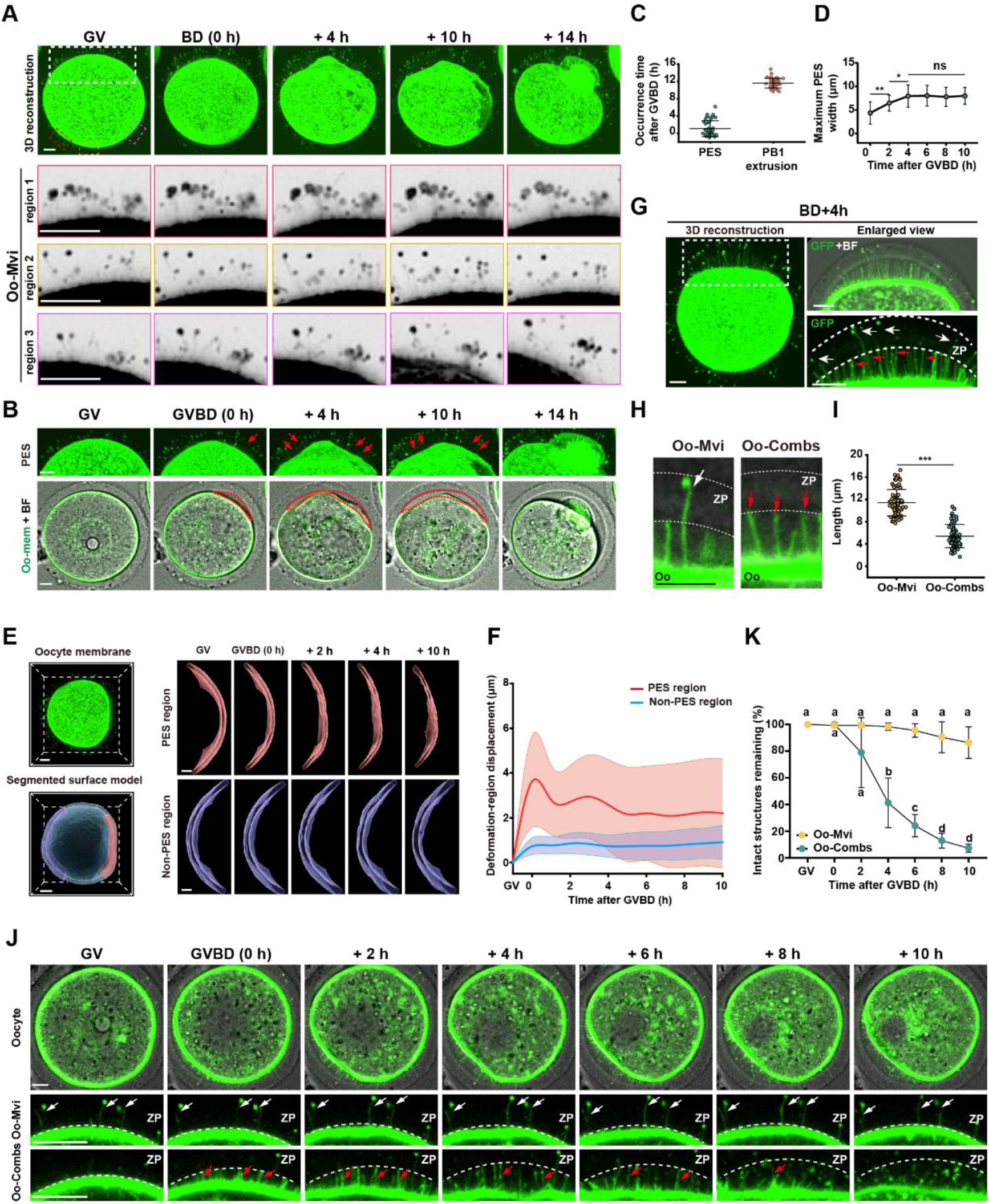
Live imaging reveals early formation of a pre-extrusion space containing transient comb-like membrane protrusions. A. Representative three-dimensional time-lapse reconstructions of the plasma membrane and associated subcellular structures in an *Oct4-CreERᵀ²;mTmG* oocyte undergoing in vitro maturation. The lower magnified time-lapse images of the regions indicated by the red, yellow and magenta boxes from the upper row, showing the stable spatial positioning of representative Oo-Mvi. Time is shown relative to GVBD (0 h). B. Representative three-dimensional time-lapse reconstructions (upper row, magnified from Fig.1A white box) and corresponding membrane-GFP and bright-field images (lower row) showing the emergence of the PES (red dashed outlines and red arrows) around GVBD and its subsequent expansion during oocyte maturation. C. Quantification of the first detection time of the PES and the timing of PB1 extrusion relative to GVBD, showing that the PES was detected approximately 10 h before PB1 extrusion (n = 43 oocytes). D. Quantification of the maximum distance between the oocyte plasma membrane and the ZP during oocyte maturation, showing that PES expansion reached a plateau by approximately 4 h after GVBD (n = 21 oocytes). E. Three-dimensional segmentation and modeling of distinct membrane regions. Left, three-dimensional reconstruction of an oocyte and schematic segmentation of the PES-associated and non-deformed regions. Right, time-lapse models of the deformed PES-associated region (PES region, red, top) and the other non-deformed region (Other region, blue, bottom), illustrating their distinct membrane dynamics. F. Quantification of membrane displacement in the PES-associated (PES region, red) and non-deformed regions (Other region, blue), showing greater and more sustained fluctuations in the PES-associated membrane (n = 10 oocytes). G. High-resolution image showing Oo-Combs (red arrows) within the PES and conventional Oo-Mvi extending into the ZP (white arrows). The ZP boundary is indicated by the dashed line. H. Magnified images comparing the morphology and length of Oo-Combs (red arrows) and Oo-Mvi (white arrows). The ZP boundary is indicated by the dashed line. I. Quantification of protrusion length, showing that Oo-Combs are significantly shorter than Oo-Mvi (n = 71 protrusions from 44 oocytes). J. Quantification of the relative abundance of Oo-Combs and Oo-Mvi during oocyte maturation. Oo-Comb abundance progressively decreases, whereas Oo-Mvi abundance remains unchanged (Oo-Combs, n = 10 oocytes; Oo-Mvi, n = 12 oocytes). For Oo-Mvi, relative abundance was normalized to the number present at GV; for Oo-Combs, which was first counted at GVBD, relative abundance was normalized to the number present at GVBD. At each subsequent time point, the numerator was the number of intact structures. K. High-resolution time-lapse images of *Oct4-CreER^T2^;mTmG* oocyte membrane during oocyte maturation, showing that Oo-Combs (bottom, red arrows) progressively extend and subsequently collapse, leading to cavity formation within the PES, whereas Oo-Mvi (middle, white arrows) remain relatively stable. The ZP boundary is indicated by the dashed line. Scale bars, 10 μm. GVBD was defined as 0 h. Data in C, D, F, I and K are shown as mean ± s.d. ns, not significant. *P < 0.05, **P < 0.01 and ***P < 0.001. Statistical significance was assessed using two-tailed unpaired Student’s *T*-tests in B and H and one-way ANOVA followed by Tukey’s multiple-comparisons test in C and I. And different letters indicate significant differences, whereas the same letters indicate no significant difference in K.

This imaging system enabled high-resolution visualization and analysis of dynamic changes in the oocyte plasma membrane and the subcellular membrane structures present on its surface throughout maturation (Fig.1A-B). Using this approach, we observed pronounced regional heterogeneity in membrane dynamics, revealing a polarized pattern of membrane behaviors across the oocyte surface (Fig.1A-B). Across most of the oocyte surface, the plasma membrane remained relatively stable, and conventional oocyte microvilli (Oo-Mvi) largely retained their density (Fig. 1A, lane 2-4 and Fig. S1D), morphology (Fig. 1A, lane 2-4 and Fig. S1E) and vesicle size (Fig. 1A, lane 2-4 and Fig. S1F) before PB1 extrusion. In striking contrast, a region of the oocyte plasma membrane underwent pronounced and spatially restricted retraction (Fig. 1B, red arrows), progressively retracting away from the zona pellucida (ZP) and generating a spatially restricted cavity (Fig. 1B, red dotted outline) between the oocyte membrane and the ZP. We termed this early, spatially restricted cavity the pre-extrusion space (PES).

This local membrane asymmetry was first detected at 1.06 ± 1.82 h relative to GVBD, 11.60 ± 1.22 h before PB1 extrusion (Fig. 1C). Localized membrane retraction progressively increased the separation between the oocyte surface and the ZP, generating a discrete cavity that reached a stable size by approximately 4 h after GVBD (Fig. 1D). To characterize the regional membrane dynamics associated with PES formation, we segmented the PES-associated deformation (red) and non-deformation stable (blue) membrane regions on the oocyte and generated 3D reconstructions by Imaris software (Fig. 1E and Movie S2). The position of the PES-associated membrane (PES region) fluctuated continuously over time, reaching a maximum displacement of 4.85 ± 2.78 μm (Fig. 1E-F, red), whereas the stable membrane (Non-PES region) remained comparatively stable, with a maximum displacement of 0.90 ± 0.48 μm (Fig. 1E-F, blue). These measurements demonstrate that PES formation arises from local spatially confined membrane dynamics rather than uniform displacement of the entire oocyte surface. Next, we reconstructed the oocyte cell body and ZP to assess volumetric dynamics (Fig. S1G). We found that the size of the oocyte decreased continuously until PB1 extrusion, while the ZP size remained unchanged during oocyte maturation (Fig. S1G-H). PES formation therefore occurs against a background of global oocyte volume reduction but is manifested as a spatially restricted separation of the oocyte membrane from a geometrically stable ZP.

High-resolution imaging further revealed that PES formation was accompanied by a distinctive membrane architecture that was not observed over the remainder of the oocyte surface. Within the developing PES, arrays of short, parallel membrane protrusions emerged from the oocyte plasma membrane and extended toward the ZP (Fig. 1G-H, red arrows). We termed these structures oocyte comb-like protrusions (Oo-Combs). In contrast to conventional Oo-Mvi (Fig. 1G-H, white arrows), Oo-Combs were organized into locally aligned arrays, projected into the widening oocyte-ZP space without penetrating the ZP (Fig. 1G-H, red arrows), and were significantly shorter than Oo-Mvi (Fig. 1G-I). Live staining with SiR-actin further revealed F-actin within Oo-Combs (Fig. S1I, white arrowheads), identifying them as actin-containing membrane protrusions associated with the locally remodeling membrane domain.

We next examined the temporal behavior of Oo-Combs during oocyte maturation. Oo-Combs initially formed and extended within the developing PES but began to collapse approximately 4 h after GVBD, coincident with the plateau in PES expansion (Fig. 1J-K, red arrows). By 10 h after GVBD, only 7.29 ± 2.26% of the initial Oo-Combs remained, leaving the PES nearly devoid of these structures and converting it into an open cavity capable of accommodating the subsequently extruded PB1 (Fig. 1J-K, red arrows). In contrast, the number of Oo-Mvi showed no significant change during oocyte maturation (Fig. 1J-K, white arrows). Together, these observations reveal an early polarized pattern of oocyte-surface behavior: a local spatially restricted domain undergoes membrane retraction, transient Oo-Comb formation and subsequent conversion into an open PES.

### The PES marks the preferred site of PB1 extrusion and correlates with PB1 size

To determine whether the PES is spatially and functionally associated with PB1 extrusion, we performed time-lapse imaging at 1-minute intervals for 1.5 hours to monitor the behaviors of PB1 extrusion (Fig. 2A). This imaging strategy allowed us to identify the site of PB1 extrusion while minimizing ambiguity introduced by post-extrusion movement of the polar body. We found that 76.29 ± 9.01% of oocytes extruded PB1 (white arrowheads) into the pre-existing PES (Fig. 2A-Bred arrowheads). This result suggests that the PES established early in maturation preferentially marks the future site of PB1 extrusion.

**Fig. 2.**
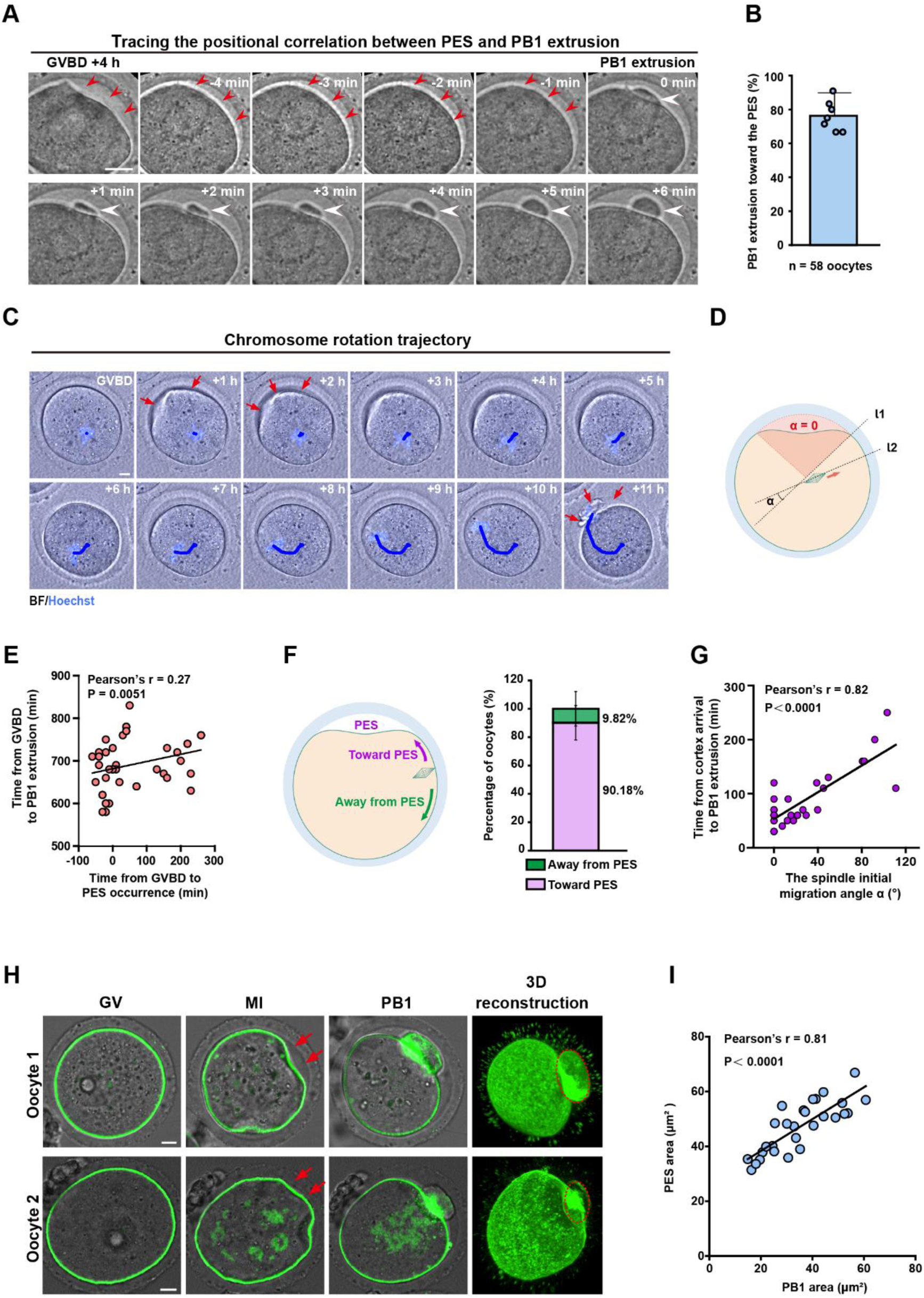
The pre-extrusion space preferentially marks the site of PB1 extrusion and scales with PB1 area. A. Representative time-lapse images acquired at 1-min intervals showing PB1 formation and extrusion within a pre-existing PES. Red arrowheads indicate the PES, and white arrowheads indicate the emerging PB1. the onset of PB1 extrusion was separately defined as 0 min. B. Quantification of the spatial relationship between the PES and PB1 extrusion. Among oocytes with a detectable PES, 76.29 ± 9.01% extruded PB1 into the pre-existing PES (n = 58 oocytes from 7 independent experiments). C. Representative time-lapse images of a live Hoechst-labelled oocyte showing the trajectory of the chromosome mass, used as a proxy for spindle position, during migration to the cortex. After reaching a cortical region outside the PES, the spindle reorients toward the PES before PB1 extrusion. The blue line denotes the chromosome trajectory, and red arrows indicate the PES. D. Schematic illustrating the measurement of the initial spindle-migration angle α relative to the PES. Vector n indicates the initial direction of spindle migration, and vector m indicates the direction from the spindle toward the PES. Direct migration toward the PES is defined as α = 0°; increasing α values indicate progressively greater deviation from the PES direction. E. Correlation between the time of PES formation and the time of PB1 extrusion. The two variables were not significantly correlated, indicating that earlier PES formation did not predict earlier PB1 extrusion (n = 43 oocytes). F. Schematic and quantification of spindle behaviors after cortical arrival (left). Among spindles that were not initially aligned with the PES (α > 0°), 90.18 ± 12.15% subsequently reoriented toward the PES before PB1 extrusion (n = 25 oocytes from 4 independent experiments). Magenta: the reorientation of spindle toward the PES; Green: the movement of spindle away from the PES. G. Correlation between the initial spindle-migration angle α and the interval from spindle arrival at the cortex to PB1 extrusion. Larger initial angular deviations were associated with longer intervals before PB1 extrusion (n = 30 oocytes). H. Representative time-lapse images of *Oct4-CreER^T2^;mTmG* oocytes with differenDt PES sizes. The oocyte in the upper panels contains a larger PES and subsequently extrudes a larger PB1 than the oocyte in the lower panels. Red arrows indicate the PES, and red dashed outlines indicate the PB1. I. Correlation between PES area and PB1 area. PES area was strongly positively correlated with the area of the subsequently extruded PB1 (n = 28 oocytes). Scale bars, 10 μm (A, C and H). Data in B and F are shown as mean ± s.d. ns, not significant. Correlations in E, G and I were assessed using two-tailed Pearson correlation analysis.

We next asked whether the PES determines the initial direction of spindle migration. To track spindle position in living oocytes, we used Hoechst-stained chromosomes as a proxy for the spindle and reconstructed their migration trajectories relative to the PES (Fig. 2C, blue trajectory line and Movie S3). We defined the initial migration angle α as the angle between the initial spindle-migration vector and the direction of the PES (Fig. 2D). Of the 30 oocytes analyzed (Fig. 2G), 13 spindles initially migrated directly toward the PES (α = 0°, red region in Fig. 2D), 14 followed an oblique trajectory relative to the PES (0° < α ≤ 90°), and 3 initially migrated away from the PES (α > 90°). These heterogeneous trajectories indicated that the initial phase of spindle migration was not consistently oriented toward the PES. Additionally, the timing of PB1 extrusion was not correlated with either the time of PES formation (Fig. 2E, Pearson’s r = 0.27) or the initial direction of spindle migration (Fig. S2D, Pearson’s r = 0.22). Thus, neither the timing of PES formation nor the initial direction of spindle migration appeared to determine the overall timing of PB1 extrusion.

We then examined spindle behaviors after the spindle reached the oocyte cortex. Among spindles that were not initially aligned with the PES (α > 0°), 90.18 ± 12.15% underwent cortical reorientation toward the PES before PB1 extrusion (Fig. 2F). Additionally, the time interval between spindle arrival at the cortex and PB1 extrusion showed a strong positive correlation with the initial spindle migration angle (α) (Fig. 2G, Pearson’s r = 0.82). Together, these observations support a two-step model in which initial spindle migration is not specified by the PES, whereas subsequent cortical repositioning preferentially converges on the PES-associated membrane domain.

Furthermore, our imaging revealed that PB1 area varied markedly among oocytes, ranging from 148.92 μm^2^ to 668.54 μm^2^ (Fig. 2H-I). In addition, we observed that oocytes with a relatively larger PES produced a correspondingly larger PB1 (red dotted line) in live-imaged *Oct4-CreER^T2^;mTmG* oocytes (Fig. 2H). Correlation analysis showed that PES size was positively correlated with PB1 size (Fig. 2I, Pearson’s r = 0.81). Thus, PES area scales with the area of the subsequently extruded PB1.

Previous studies have shown that a polarized increase in cortical F-actin thickness contributes to oocyte shape remodeling and spindle positioning through Arp2/3-dependent actin dynamics^23, 24^. To investigate whether PES formation is driven by a polarized increase in cortical thickness, we measured the cortex thickness in both deformation and non-deformation areas during maturation. We found that cortical thickness in the deformation area increased at the MI stage (GVBD + 6 h), which occurs after PES formation (1.06 ± 1.82 h relative to GVBD) (Fig. S2A-C). Thus, PES formation precedes the establishment of the canonical actin-enriched cortical domain, arguing against late cortical F-actin thickening as the initiating cause of PES formation. These results identify the PES as an early spatial asymmetry that prefigures the later cortical and spindle positioning associated with PB1 extrusion.

### ERK-mediated suppression of ZO-1 phase separation promotes its local dissociation before PES formation

Because PES formation begins with localized retraction of the oocyte plasma membrane away from the ZP, we reasoned that this process might involve remodeling of membrane-associated cortical scaffolds. Although canonical mature tight junctions are not present at the oocyte-GC interface, ZO-1 (TJP1) as a scaffold protein has been detected in oocytes and at this interface as part of a broader junction-related cortical protein network^34^. We therefore investigated whether ZO-1 is associated with PES formation. We first examined ZO-1 localization in *Oct4-CreER^T2^;mTmG* oocytes by immunofluorescent staining and found that ZO-1 formed discrete puncta immediately beneath the oocyte plasma membrane (Fig. 3A-B). Fluorescence intensity analysis showed that, at 2 h after GVBD, the PES-associated retracted membrane (PES, red box) exhibited fewer and more dispersed ZO-1 puncta than the non-deformed membrane region (Non-PES, blue box) (Fig. 3C-D). In addition, we found that overall ZO-1 immunofluorescence intensity progressively decreased during in vitro maturation (Fig. 3C and E). These data associate local loss of membrane-associated ZO-1 with PES formation.

**Fig. 3.**
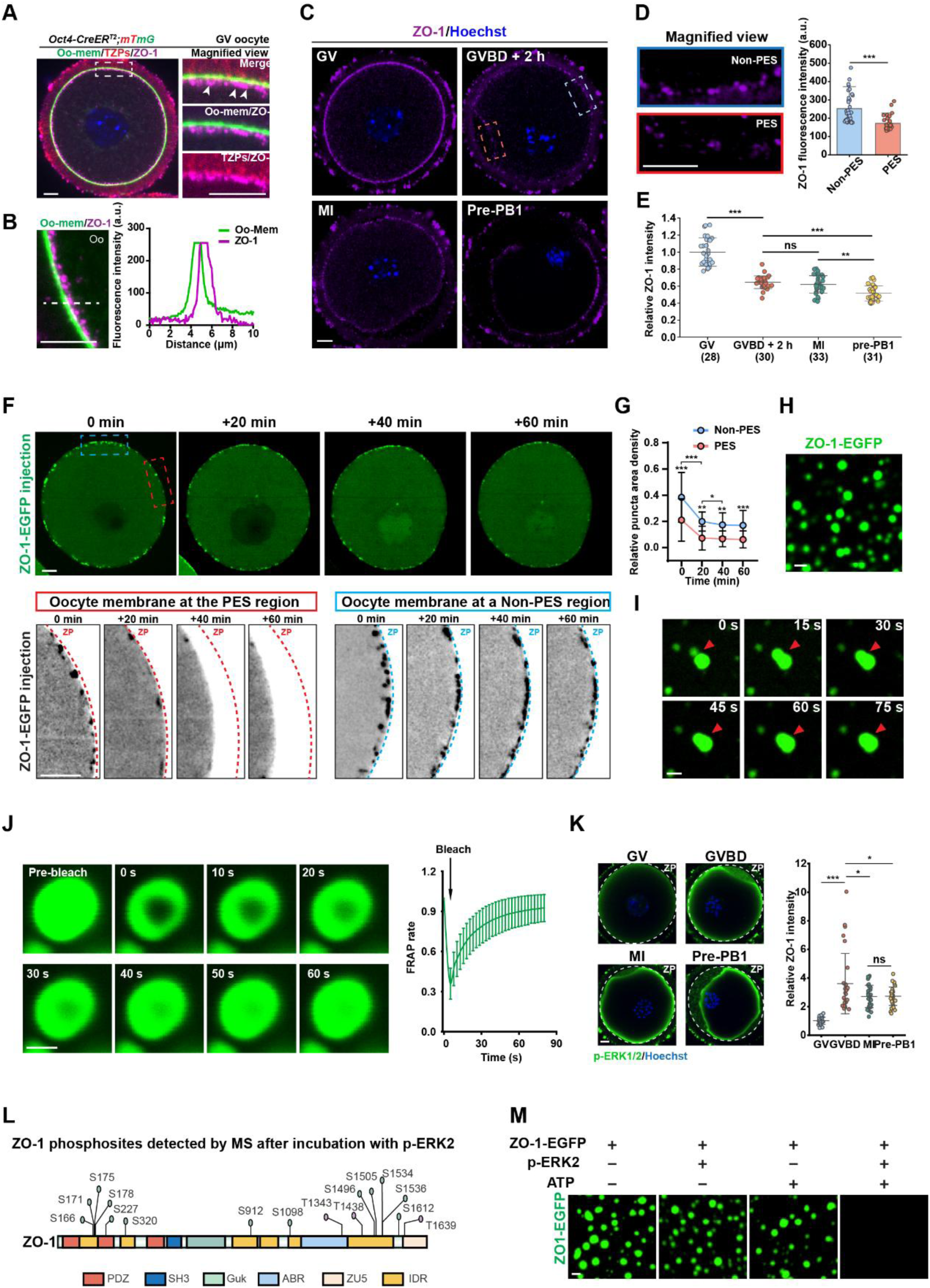
ZO-1 depletion precedes PES formation, and ERK2 phosphorylation suppresses ZO-1 phase separation in vitro. A. Representative images of *Oct4-CreER^T2^;mTmG* oocytes stained for ZO-1. Magnification of the boxed region shows ZO-1 puncta immediately beneath the oocyte plasma membrane. White arrowheads indicate ZO-1 puncta. B. Line-scan fluorescence intensity profiles along the dashed line in A, showing the submembranous localization of ZO-1 puncta relative to the membrane-GFP signal. C. Representative images of ZO-1 localization at the indicated stages of oocyte maturation. At 2 h after GVBD, membrane retraction in the PES-associated region was accompanied by local dispersal and loss of ZO-1 puncta (red box), whereas the non-PES region retained a stronger punctate ZO-1 signal (blue box). D. Magnification of the regions indicated in C (left) and paired quantification of ZO-1 fluorescence intensity at 2 h after GVBD (right), showing lower ZO-1 intensity in the PES-associated region than in the non-PES region (n = 36 oocytes). E. Quantification of overall cortical ZO-1 fluorescence intensity at the indicated stages, showing a progressive decrease during oocyte maturation (n = 28, 30, 33 and 31 oocytes, respectively). F. Representative time-lapse images of an oocyte microinjected with purified ZO-1-EGFP. At the beginning of imaging, ZO-1-EGFP puncta were less abundant in the membrane region that subsequently formed the PES (PES region; red box) than in the non-PES region (non-PES region, blue box). ZO-1-EGFP density declined further before detectable membrane retraction, and puncta were nearly absent from the PES-associated membrane by 60 min. The dashed line indicates the inner boundary of the ZP. Dashed line: zona pellucida. The time of capturing first picture of oocyte was defined as 0 min. G. Quantification of relative ZO-1-EGFP puncta area density in the regions shown in F. Puncta density gradually decreased in both regions but remained consistently lower in the prospective PES region than in the non-PES region throughout the imaging period (n = 11 oocytes). H. Representative fluorescence image of condensates spontaneously formed by purified ZO-1-EGFP in vitro. I. Time-lapse images showing fusion of two ZO-1-EGFP condensates. Red arrowheads indicate the fusing condensates. J. Representative images and quantification of fluorescence recovery after photobleaching (FRAP) of ZO-1-EGFP condensates. Fluorescence recovered to 92.5 ± 10.1% within 90 s, indicating rapid molecular exchange within the condensates (n = 12 condensates from 3 independent assays). K. Representative immunofluorescence images and quantification of cortical phosphorylated ERK1/2 (p-ERK1/2) intensity at the indicated stages. Cortical p-ERK1/2 increased markedly at GVBD and subsequently declined during oocyte maturation (n = 27, 29, 31 and 31 oocytes, respectively). Hoechst marks chromatin, and the dashed outline indicates the ZP. Relative intensity was normalized to the average intensity at GV. L. Mapping of ZO-1 phosphorylation sites identified by mass spectrometry following incubation with active p-ERK2 and ATP in vitro. Mass spectrometry identified 16 phosphorylated residues, 12 of which were located within intrinsically disordered regions. M. Representative images of in vitro phase-separation assays showing that addition of active p-ERK2 and ATP markedly suppressed ZO-1-EGFP droplet formation. The four assay conditions were ZO-1-EGFP alone, ZO-1-EGFP with p-ERK2, ZO-1-EGFP with ATP, and ZO-1-EGFP with both p-ERK2 and ATP. Scale bars, 10 μm (C, D, F and K), 2 μm (H, I and M) and 1 μm (J). Data in D, G, J and K are shown as mean ± s.d. ns, not significant. *P < 0.05, **P < 0.01 and ***P < 0.001. Statistical significance was assessed using a two-tailed paired Student’s t-test in D and one-way ANOVA followed by Tukey’s multiple-comparisons test in E and K.

To further define the temporal relationship between ZO-1 dynamics and PES formation, we expressed and purified the full-length mouse ZO-1 protein fused to C-terminal mEGFP and Flag tag (ZO-1-EGFP) using a eukaryotic insect-cell expression system (Fig. S3B, left). As the negative control, mEGFP protein was injected into oocytes, producing a uniform cytoplasmic fluorescence signal (Fig. S3A). Next, we injected ZO-1-EGFP into GV-stage oocytes and monitored its dynamics by high resolution live-cell imaging during culture. Consistent with the immunofluorescence data, ZO-1-EGFP formed discrete punctate assemblies that were non-uniformly distributed along the oocyte cortex at the beginning of imaging before GVBD (Fig. 3F, 0 min). Quantification revealed that the membrane region that would subsequently give rise to the PES already exhibited a markedly lower ZO-1-EGFP puncta density at 0 min than the remainder of the oocyte membrane (Fig. 3F and S3C). Live-cell imaging showed ZO-1-EGFP density in the prospective PES region declined further by 20 min, before detectable membrane deformation (Fig. 3F-G). The same region subsequently retracted from the ZP between 40 and 60 min and developed into the PES (Fig. 3F-G). By 60 min of live-cell imaging, ZO-1-EGFP puncta were nearly absent from this retracted membrane region (Fig. 3F-G). More broadly, ZO-1-EGFP density declined progressively in both regions throughout imaging, while remaining consistently lower in the prospective PES region than in the non-PES region (Fig. 3F-G). Thus, PES formation occurs within a ZO-1-low membrane domain and is preceded by further ZO-1 depletion before detectable membrane retraction.

ZO-1 contains intrinsically disordered linker regions and a conserved PDZ3-SH3-GuK domain, structural features that support multivalent interactions^40^. We therefore hypothesized that its phase behavior might regulate its local dissociation from the oocyte membrane. Indeed, purified ZO-1-EGFP spontaneously formed liquid-like droplets that fused over time (Fig. 3H-I). To further test whether these condensates were formed by liquid-liquid phase separation (LLPS), we performed FRAP analysis showing the fluorescence signal within photobleached regions in ZO-1-EGFP droplets recovered to 92.5 ± 10.1% within 90 s (Fig. 3J), indicating rapid molecular exchange within the condensates. Together, droplet fusion and rapid fluorescence recovery support the formation of dynamic ZO-1 condensates through LLPS in vitro.

The phase behavior of human ZO-1 has been reported to be regulated by phosphorylation in a spatially controlled manner^35^. In addition, our immunofluorescence analyses showed the cortical p-ERK1/2 signal increased markedly at GVBD, reached its highest level during early maturation and subsequently declined after MI, temporally coinciding with PES formation (Fig. 3K). To determine whether active ERK directly phosphorylates ZO-1 and alters its phase behavior, we purified active phosphorylated mouse ERK2 carrying a C-terminal Strep tag (p-ERK2) by co-expressing it with His-MEK1 in an *E. coli* system (Fig. S3B, right), and then incubated it with ZO-1 protein in vitro. Mass spectrometry identified 16 ZO-1 residues directly phosphorylated by p-ERK2 in vitro, 12 of which were located within intrinsically disordered linker regions, which related to LLPS mediated (Fig. 3L and Table S1). Importantly, addition of p-ERK2 and ATP triggered the dissolution of pre-formed ZO-1-EGFP condensates in vitro (Fig. 3M). These results show that ERK2 phosphorylation suppresses ZO-1 phase separation in vitro, providing a mechanism consistent with local ZO-1 depletion from the prospective PES-associated membrane. Together, these findings support a model in which ERK-dependent phosphorylation suppresses ZO-1 phase separation and may thereby promote the local loss of membrane-associated ZO-1 before PES formation.

### ERK-dependent ZO-1 remodeling supports PES establishment and PB1 extrusion

To test whether the ERK-dependent ZO-1 remodeling identified above operates in oocytes, we treated oocytes with the MEK1/2 inhibitor U0126 to suppress ERK1/2 activation^41^ and examined ZO-1 localization at 2 h after GVBD. Immunofluorescence images showed that, whereas control oocytes exhibited local depletion of ZO-1 from the PES-associated membrane, U0126-treated oocytes retained strong ZO-1 punctate signals throughout the cortex and lacked the localized ZO-1 loss that normally accompanied membrane retraction (Fig. 4A-C). Together with the in vitro phosphorylation and phase-separation results, these findings support a model in which ERK-dependent phosphorylation promotes ZO-1 dissociation from the oocyte membrane during early maturation.

**Fig. 4.**
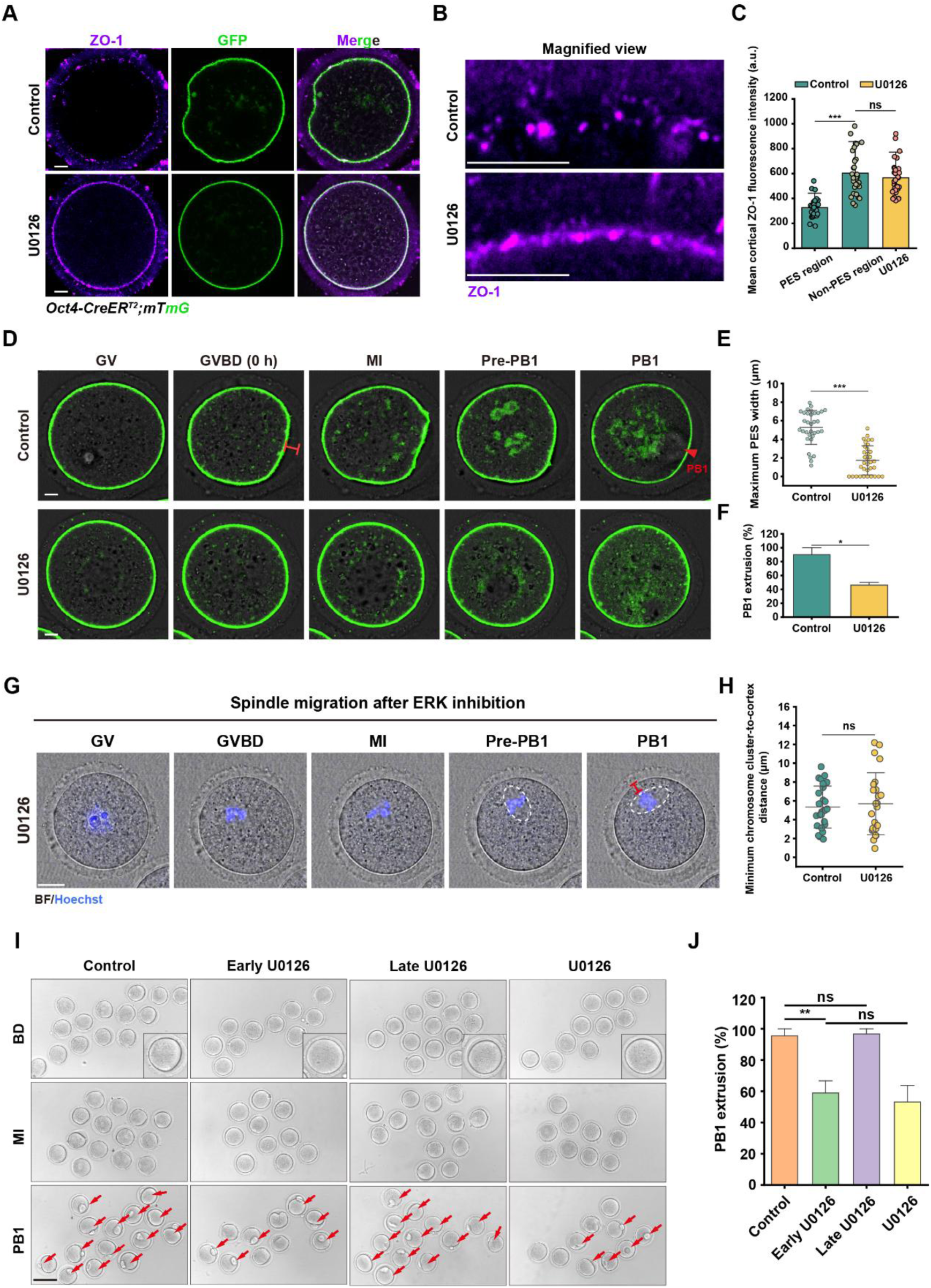
MEK–ERK inhibition disrupts ZO-1 remodeling, PES formation and PB1 extrusion. A. Representative ZO-1 immunofluorescence images of control and U0126-treated *Oct4-CreER^T2^;mTmG* oocytes at 2 h after GVBD. U0126-treated oocytes retain strong ZO-1 punctate signals throughout the cortex, whereas control oocytes exhibit an overall reduction in cortical ZO-1 together with further local depletion at the PES-associated membrane. B. Magnified views of the boxed regions in A. Control oocytes show localized loss of ZO-1 puncta from the PES-associated membrane, whereas U0126-treated oocytes retain ZO-1 puncta and fail to establish a discrete ZO-1-low membrane domain. C. Quantification of ZO-1 fluorescence intensity in the PES-associated and non-PES membrane regions of control oocytes and across the cortical membrane of U0126-treated oocytes. In control oocytes, ZO-1 intensity was lower in the PES-associated region than in the non-PES region, whereas U0126-treated oocytes retained a high cortical ZO-1 signal without detectable regional depletion (n = 32, 32 and 31 oocytes, respectively). D. Representative time-lapse images of control and U0126-treated *Oct4-CreER^T2^;mTmG* oocytes during maturation. Control oocytes form a prominent PES and subsequently extrude PB1, whereas U0126-treated oocytes exhibit impaired PES formation and fail to extrude PB1 in the representative example. Red arrowhead: PB1. E. Quantification of the maximum distance between the oocyte plasma membrane and the ZP. U0126 treatment results in absent or narrowed PES formation and significantly reduces the maximum membrane-ZP distance compared with controls (control, n = 33 oocytes; U0126, n = 34 oocytes). F. Quantification of PB1 extrusion showing that U0126 treatment reduces the PB1 extrusion rate (n = 3 independent experiments). G. Representative time-lapse images of Hoechst-labelled oocytes showing migration of the chromosome-associated spindle toward the cortex in control and U0126-treated oocytes. Dashed ellipses: the chromosome-associated spindle position; Red line: the minimum distance between the chromosomes and the plasma membrane. H. Quantification of the minimum chromosome-to-plasma membrane distance, showing no significant difference between control and U0126-treated oocytes (control, n = 26 oocytes; U0126, n = 24 oocytes). I. Representative images of oocytes subjected to the indicated U0126 treatment regimens, showing the corresponding effects on PES formation and PB1 extrusion. Continuous and early U0126 treatment impair PES formation and PB1 extrusion, whereas oocytes treated with U0126 after the early establishment window retain a visible PES and extrude PB1. J. Quantification of PB1 extrusion following the indicated U0126 treatment regimens. Continuous and early U0126 treatment significantly reduce PB1 extrusion, whereas the extrusion rate following late U0126 treatment is comparable to that in untreated controls (n = 3 independent experiments per group). Scale bars, 10 μm (A, B, D and G) and 100 μm (I). Data in C, E, F, H and J are shown as mean ± s.d. ns, not significant. *P < 0.05, **P < 0.01 and ***P < 0.001. Statistical significance was assessed using two-tailed unpaired Student’s t-tests in C, E and F.

Next, live-cell imaging of *Oct4-CreER^T2^;mTmG* oocyte maturation showed that U0126 treatment impaired localized membrane dynamics and PES formation (Fig. 4D and Movie S4). Among U0126-treated oocytes, 29.41% completely lacked a detectable PES, whereas the remaining 70.59% formed a smaller PES than control oocytes (Fig. 4D-E). Accordingly, the maximum distance between the oocyte plasma membrane and the ZP decreased from 5.28 ± 1.82 μm in controls to 2.45 ± 1.29 μm following U0126 treatment (Fig. 4D-E). Consistent with impaired PES formation, PB1 extrusion decreased from 90.21 ± 11.34% in control oocytes to 46.10 ± 3.61% in U0126-treated oocytes (Fig. 4F).

To assess whether the reduction in PB1 extrusion could be explained by gross defects in spindle positioning or organization, we tracked Hoechst-labelled chromosomes as a proxy for spindle migration in living oocytes. The chromosome-associated spindle reached the cortex in U0126-treated oocytes, and the final spindle-to-cortex distance was comparable between control and U0126-treated groups (Fig. 4G-H, 5.34 ± 2.22 μm vs. 5.69 ± 3.29 μm). Immunofluorescence staining for α-tubulin further revealed no detectable alteration in gross spindle morphology following U0126 treatment (Fig. S4A-B). Thus, the reduction in PB1 extrusion following ERK inhibition was associated with defective PES formation despite preserved cortical spindle positioning and gross spindle organization.

We next asked whether ERK-dependent PES establishment is confined to an early maturation window or can be deferred to later stages. Oocytes were treated with U0126 during the first 6 h of in vitro maturation, ending at approximately 4 h after GVBD, and were then washed and cultured in inhibitor-free M2 medium (Early U0126; Fig. S4C, Lane2). Similar to continuous U0126 exposure (U0126; Fig. S4C, Lane3), early U0126 treatment in persistent PES defects and reduced PB1 extrusion, despite subsequent inhibitor removal (Fig. 4I-J and Fig. S4D). By contrast, when oocytes were cultured in M2 medium for the first 6 h and then exposed to U0126 (Late U0126; Fig. S4C, Lane4), after the PES had normally reached a stable state, both PES area and PB1 extrusion were comparable to those in untreated controls (Fig. 4I-J and Fig. S4D). Together, these findings define a critical early ERK-dependent window for PES establishment. Late U0126 treatment did not detectably affect the maintenance of the established PES. Conversely, when PES formation is disrupted during this early window, subsequent removal of ERK inhibition is insufficient to establish a normal PES later in maturation.

### ERK activation restores PES formation and partially rescues PB1 extrusion in aged oocytes

We next asked whether defective early membrane remodeling contributes to the age-associated decline in PB1 extrusion. To examine PES formation during reproductive ageing, we tracked plasma membrane dynamics in oocytes isolated from young (2-month-old) and aged (12-month-old) mice. High-resolution time-lapse imaging revealed that young oocytes typically formed a PES accompanied by the appearance and subsequent collapse of Oo-Combs, whereas a subset of aged oocytes failed to establish a detectable PES (Fig. 5A-B and Movie S5). The proportion of oocytes lacking a PES increased from 5.51 ± 6.47% in young mice to 21.39 ± 7.84% in aged mice (Fig. 5B). Among oocytes that formed a detectable PES, PES area was smaller in aged oocytes than in young controls (Fig. 5F, young vs. aged: 372.55 ± 125.70 μm² vs. 162.65 ± 89.96 μm²). Consistent with these PES defects, PB1 extrusion decreased from 87.25 ± 3.80% in young oocytes to 51.26 ± 13.90% in aged oocytes (Fig. 5C). Thus, defective PES formation is an age-associated oocyte phenotype that accompanies reduced PB1 extrusion.

**Fig. 5.**
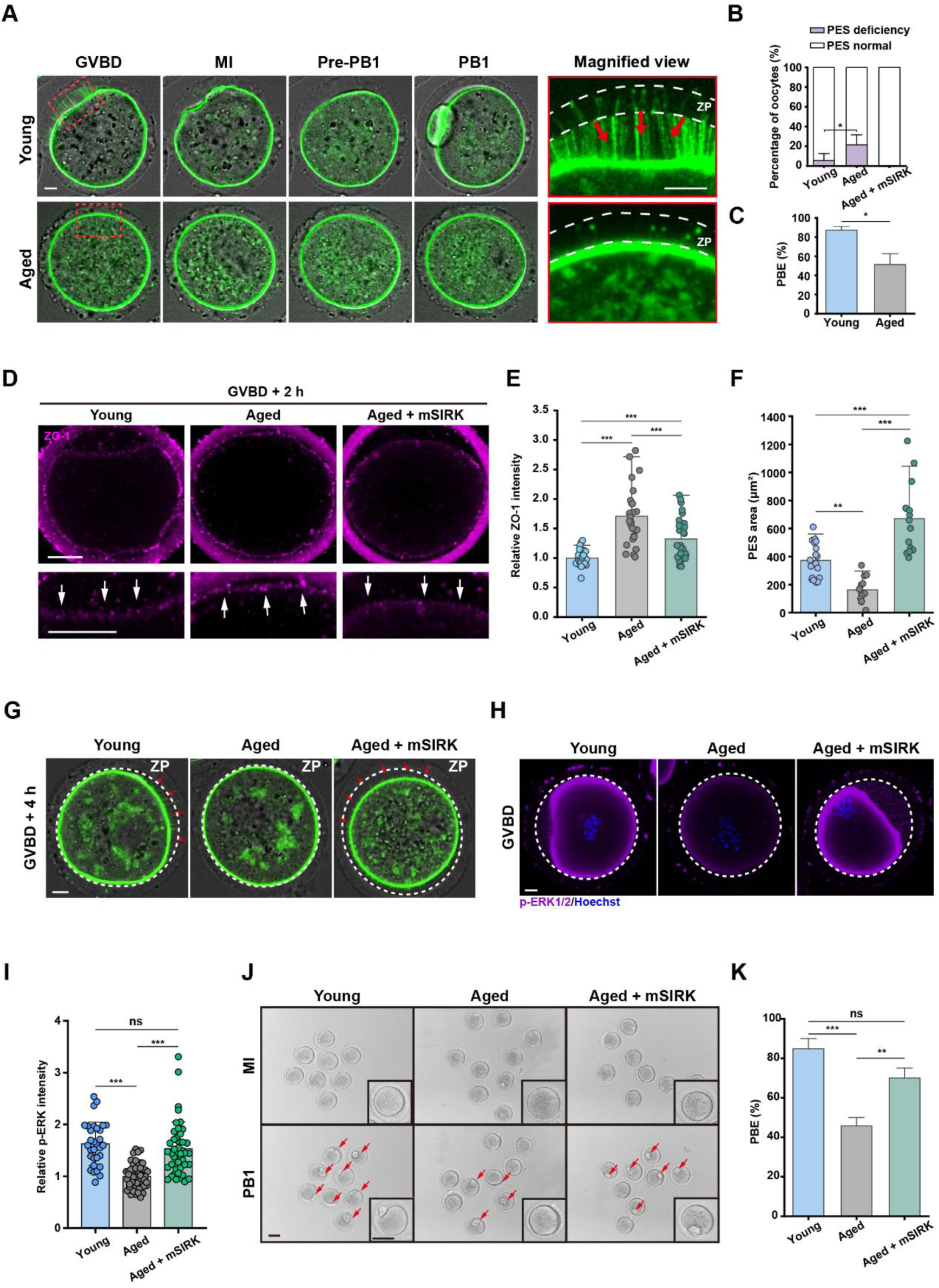
Enhanced ERK activity restores PES formation and partially rescues PB1 extrusion in aged oocytes. A. Representative time-lapse images showing impaired PES formation, reduced PB1 extrusion and loss of Oo-Combs (red arrows) in aged oocytes compared with young oocytes. Dashed line: zona pellucida. Magnified views of the regions outlined by the red dashed boxes. B. Quantification of PES occurrence shows that PES is increasingly absent in old oocytes (n = 3 independent experiments). C. Quantification of oocyte maturation indicates compromised PB1 extrusion in old oocytes (n = 3). D. Representative immunofluorescence images of oocytes at 2 h after GVBD stained with an anti-ZO-1 antibody show that cortical ZO-1 is retained in the PES-associated region (white arrows) in old oocytes and is reduced following mSIRK treatment. Lower panels show magnified views of the corresponding cortical regions in the upper panels. E. Quantification of ZO-1 fluorescence intensity shows that mSIRK treatment decreases the elevated ZO-1 signal in old oocytes (n = 41, 25, 29 respectively). Relative intensity was normalized to the average intensity at GV. F. Quantification of PES occurrence shows that PES formation is reduced in old oocytes and promoted by mSIRK treatment (n = 18, 13, 15 respectively). G. mSIRK treatment restores detectable PES formation (red arrows). Dashed ellipse: zona pellucida. Red marks indicate PES. H. Representative p-ERK immunofluorescence images of young, aged and mSIRK-treated oocytes at 2 h after GVBD. Cortical p-ERK1/2 signal is lower in aged oocytes than in young oocytes and is increased following mSIRK treatment. The dashed outline indicates the ZP. I. Quantification of p-ERK fluorescence intensity in cortex of young, aged and mSIRK-treated oocytes. In aged oocytes, p-ERK intensity was lower than young, whereas mSIRK treatment increased cortical p-ERK1/2 signal in aged oocytes (n =31, 52, 43 respectively). J. Representative images showing increased PB1 extrusion (red arrows) in mSIRK-treated aged oocytes compared with untreated aged oocytes. K. Quantification of PB1 extrusion in young, aged and mSIRK-treated aged oocytes, showing partial recovery following mSIRK treatment (n = 3 independent experiments). Scale bars, 10 μm (A, D, G and H). Data are shown as mean ± s.d.; ns, not significant; *P < 0.05, **P < 0.01, ***P < 0.001.

We next examined whether ERK-dependent ZO-1 remodeling was altered in aged oocytes. At 2 h after GVBD, during PES formation, cortical ZO-1 intensity was higher in aged oocytes than in young controls (Fig. 5D-E, young vs. aged: 1.00 ± 0.11 vs. 1.71 ± 0.50). Conversely, p-ERK1/2 intensity was lower in aged oocytes than in young controls (Fig. 5H-I, young vs. aged: 1.63 ± 0.41 vs. 1.00 ± 0.23). These results are consistent with the effects of U0126 and associate impaired ERK-dependent ZO-1 remodeling with defective PES formation in aged oocytes.

To test whether this defect was reversible, we treated aged oocytes with the ERK activator mSIRK^42^ and confirmed that p-ERK1/2 intensity increased following treatment (Fig. 5H-I, aged vs. aged + mSIRK: 1.00 ± 0.23 vs. 1.53 ± 0.51). At 2 h after GVBD, mSIRK treatment reduced cortical ZO-1 intensity from 1.71 ± 0.50 in untreated aged oocytes to 1.32 ± 0.36 (Fig. 5D-E). Moreover, the proportion of aged oocytes lacking a detectable PES decreased from 21.39 ± 7.84% to 0%, indicating restoration of PES formation across the treated population (Fig. 5B). In addition, imaging of *Oct4-CreER^T2^;mTmG* oocytes showed that PES area also increased from 162.65 ± 89.96 μm² in untreated aged oocytes to 670.76 ± 248.94 μm² following mSIRK treatment (Fig. 5F-G, red arrows). Thus, mSIRK treatment enhanced ERK activity, promoted ZO-1 remodeling and restored PES establishment in aged oocytes.

Importantly, mSIRK treatment increased PB1 extrusion from 45.77 ± 3.75% in untreated aged oocytes to 70.00 ± 8.66% (Fig. 5J-K). Notably, the improvement in PB1 extrusion closely matched the proportion of aged oocytes that initially failed to form a PES (24.23% vs. 21.39%). This correspondence is consistent with mSIRK rescuing PB1 extrusion primarily in the subset of aged oocytes with defective ERK-ZO-1-dependent PES formation. Together, these findings indicate that impaired ERK-ZO-1 remodeling contributes to a component of age-associated oocyte maturation failure, suggesting that the ERK-ZO-1 pathway may represent a potential target for improving the in vitro maturation of oocytes from aged mouse oocytes.

## Discussion

Mammalian oocyte maturation requires not only the establishment of polarity and execution of highly asymmetric cytokinesis, but also adaptation to the physical confinement imposed by the surrounding ZP^1, 13^. Although extensive work has defined the contributions of actin remodeling, spindle migration and cortical mechanics to PB1 extrusion^5, 11, 23^, how the oocyte plasma membrane spatially reorganized in preparation for this process has remained unclear. Here, using a genetically encoded membrane reporter in mouse oocytes combined with high-resolution live imaging, we systematically characterized oocyte membrane regional heterogeneity dynamics during maturation. Around GVBD, a discrete region of the oocyte plasma membrane retracts from the ZP to form the PES, within which transient Oo-Combs subsequently collapse to generate an open cavity before PB1 extrusion. The PES preferentially marks the future site of PB1 extrusion, and its area is strongly correlated with that of the subsequently extruded PB1. Although the PES occupies the same oocyte–ZP compartment as the mature perivitelline space, its early, localized and dynamically generated nature distinguishes it from the conventionally recognized perivitelline space and suggests that it may contribute to, rather than simply reflect, the later perivitelline architecture^43, 44^. These findings identify PES formation as an early membrane pre-remodeling event that establishes a spatially permissive geometry before PB1 extrusion under ZP confinement (Fig. S5).

A defining feature of PES formation is that membrane remodeling is regional rather than global. The coexistence of a largely stable surface that retains Oo-Mvi with a discrete, highly dynamic domain undergoing membrane retraction and Oo-Comb turnover represents an early form of surface symmetry breaking during oocyte maturation. This spatial partitioning may be functionally advantageous, allowing the oocyte to preserve membrane organization across most of its surface while selectively generating a deformable region for PB1 extrusion. Under the mechanical confinement imposed by the ZP, localized membrane disengagement may therefore provide an efficient strategy for creating the required spatial clearance without broadly disrupting the oocyte-ZP interface. More broadly, these findings suggest that regional regulation of membrane-cortex organization can preconfigure cell geometry before asymmetric division.

Previous studies have placed the emergence of oocyte cortical polarity in mid-to-late meiosis I, when Mos-MAPK dependent cortical F-actin thickening and myosin-II exclusion soften the cortex and establish a permissive mechanical window for spindle migration^23, 24^. Our findings extend the onset of detectable polarity to a substantially earlier stage. ZO-1 is locally depleted from the oocyte cortex before detectable membrane retraction, and the PES becomes morphologically apparent around GVBD, several hours before the polarized increase in cortical F-actin thickness at approximately 6 h after GVBD. These observations indicate that oocyte surface polarity is initiated at the molecular level and becomes morphologically manifest through PES formation around GVBD, well before the establishment of the canonical actin-enriched cortical domain. Thus, our findings challenge the prevailing view that the oocyte lacks detectable polarity during early meiotic maturation^45, 46^ and instead reveal a temporal hierarchy of polarization, beginning with asymmetric ZO-1 organization, followed by localized membrane retraction and PES formation, and culminating in later actin-dependent cortical polarization. Although the temporal sequence does not establish that the PES directly induces cortical F-actin remodeling, the preferential reorientation of cortically positioned spindles toward the PES suggests that this early polarized membrane domain may provide a spatial landmark upon which later cortical remodeling and spindle positioning converge.

ZO-1 is best known as a scaffold at epithelial tight junctions, where it connects transmembrane adhesion complexes to cortical actin and contributes to the spatial and mechanical organization of cell-cell interfaces^32, 33, 47^. However, mammalian oocytes do not form canonical epithelial tight junctions with the surrounding granulosa cells^34^. Notably, we found that ZO-1 is expressed in oocytes and exhibits a punctate, spatially heterogeneous cortical distribution, rather than the continuous junctional pattern characteristic of epithelial cells. This atypical localization suggests that ZO-1 performs a noncanonical scaffolding function at the oocyte cortex. ZO-1 distribution is already reduced in the prospective PES region and declines further before local membrane retraction. In vitro, purified ZO-1 forms dynamic condensates, whereas direct phosphorylation by active ERK2 suppresses its phase separation and dissolves preformed condensates. Consistent with these biochemical findings, inhibition of ERK activation with U0126 prevents the localized loss of cortical ZO-1, impairs PES formation and reduces PB1 extrusion. These effects occur without detectable disruption of cortical spindle positioning or gross spindle organization, indicating that the reduction in PB1 extrusion is not readily explained by overt spindle defects. Together, these observations support a model in which ERK-dependent phosphorylation suppresses ZO-1 condensation and promotes localized removal of the cortical scaffold, thereby permitting membrane retraction and PES formation. Although the purified-protein experiments establish the intrinsic capacity of ZO-1 to undergo phosphorylation-sensitive phase separation, direct demonstration of endogenous ZO-1 phase behavior at the oocyte cortex will be required to determine whether bona fide LLPS occurs in vivo. Nevertheless, our findings indicate that PES formation is an actively regulated membrane-remodeling process and suggest that the phase behavior of junction-associated scaffolds may be repurposed to generate subcellular asymmetry structure in oocyte.

Age-associated deterioration in oocyte quality and IVM competence remains a major obstacle to female fertility^48, 49^. Previous studies have identified chromosomal abnormalities in aged oocytes^50–52^ as well as broader defects in mitochondrial and protein homeostasis^53, 54^. However, age-related defects in oocyte membrane remodeling remain poorly understood. We found that a subset of aged oocytes failed to undergo localized membrane retraction and consequently exhibited defective PES formation. This phenotype was accompanied by reduced cortical p-ERK1/2 and increased retention of ZO-1 at the oocyte membrane, recapitulating key features of pharmacological ERK inhibition. Treatment with mSIRK increased p-ERK1/2, reduced cortical ZO-1 retention and eliminated detectable PES absence in the analyzed population of aged oocytes. However, PB1 extrusion was only partially restored. Notably, the increase in PB1 extrusion closely corresponded to the proportion of aged oocytes that lacked a PES, consistent with preferential rescue of a PES-defective subpopulation. This population-level correspondence does not establish that the same individual oocytes were rescued, but it supports the conclusion that defective ERK-ZO-1-dependent PES formation accounts for a defined component of age-associated maturation failure. Further studies should also determine how reproductive ageing affects the mechanical properties and molecular composition of the oocyte surface and whether this pathway can be safely manipulated in human oocytes. Together, these findings indicate that impaired ERK-ZO-1 remodeling contributes to age-associated oocyte maturation failure in a subset of oocytes, suggesting that the ERK-ZO-1 pathway may represent a potential target for improving IVM outcomes in oocytes from aged females.

Overall, we use high-resolution live oocyte imaging system to identify PES formation as a membrane pre-remodeling event that begins hours before PB1 extrusion. Our research demonstrates that local depletion of cortical ZO-1 precedes membrane retraction, while ERK directly phosphorylates ZO-1 and suppresses its phase separation in vitro. Inhibition of MEK-ERK signaling prevents ZO-1 remodeling, disrupts PES formation and reduces PB1 extrusion. In aged oocytes, reduced ERK activation and cortical retention of ZO-1 are associated with defective PES formation, whereas pharmacological enhancement of ERK activity restores PES establishment and partially rescues PB1 extrusion. Together, these findings establish membrane pre-remodeling as an early architectural step in PB1 extrusion and identify defective ERK-dependent ZO-1 remodeling as one component of oocyte ageing.

## Materials and methods

### Mice

Animal experiments were carried out in compliance with the laboratory animal care and use guidelines of China Agricultural University and were approved by the Animal Welfare and Ethics Committee (approval no. AW72012202-3-1).

ICR mice were obtained from the Laboratory Animal Center of the Institute of Genetics, Beijing, China. *Oct4-CreER^T2^;mTmG* mice were produced by crossing *Oct4-CreER^T2^* mice (gifted from Dr. Ping Zheng at Yunnan Key Laboratory of Animal Reproduction, Kunming Institute of Zoology, Chinese Academy of Sciences, Kunming, China) with *mTmG* reporter mice (007576, Jackson Laboratory), both of which were generated as previously reported. Tamoxifen (TAM; Sigma-Aldrich, 75648) was prepared by dissolving it in 95% ethanol at 100 mg/ml, followed by dilution in corn oil (Sigma-Aldrich, C8267) to 20 mg/ml. For oocyte membrane labeling, *Oct4-CreER^T2^;mTmG* females received intraperitoneal injections of TAM at 100 mg/kg body weight at PD 10.

All mice were maintained under specific pathogen-free (SPF) conditions with a controlled temperature of 24-26 °C, humidity of 40-70%, and a 16/8 h light/dark cycle, with food and water provided ad libitum.

### Oocyte collection and culture

Ovaries were collected from 6-8-week-old and 12-month-old (aged) female mice. Fully grown oocytes were isolated by puncturing antral follicles and maintained in M2 medium (M1250; AIBI Bio-technology Co., Ltd., Nanjing, China) supplemented with 2.5 μM milrinone to maintain prophase I arrest during collection. To induce meiotic resumption, milrinone was removed by washing the oocytes in milrinone-free M2 medium, after which the oocytes were cultured at 37 °C and 5% CO_2_.

Oocytes were identified and examined at 0, 2, 6, 10 and 12 h after milrinone removal, corresponding to the GV, BD, MI, pre-PB1 and PB1 extruded stages, respectively.

### Immunofluorescence staining

Oocytes were sequentially fixed in 2% paraformaldehyde in PBS for 5 min and then in 4% paraformaldehyde in PBS for 10 min at room temperature to preserve oocyte morphology. After fixation, the oocytes were washed with PBS and permeabilized in PBST buffer (PBS containing 0.1% Triton X-100) for 15 min at room temperature. The oocytes were then blocked in PBS containing 3% BSA for 1 h at room temperature and incubated overnight at 4 °C with primary antibodies diluted in PBS containing 0.1% BSA. After three washes with PBS, the oocytes were incubated with the appropriate secondary antibodies for 1.5 h at room temperature. The oocytes were subsequently washed three times with PBS, stained with Hoechst 33342 (Beyotime, C1025, 1:100) for 3 min at room temperature and washed again with PBS. For imaging, the oocytes were placed in a 20-μl drop of PBS covered with paraffin oil in a 35-mm glass-bottom dish fitted with a No. 1.0 coverslip.

Primary antibodies included mouse anti-ZO-1 (Invitrogen, 33-9100, 1:100) and rabbit anti-p-ERK (Abclonal, AP0234, 1:100). Secondary antibodies included Alexa Fluor 488- or 647-conjugated, highly cross-adsorbed donkey anti-rabbit IgG (Invitrogen A21202, 1:100) and donkey anti-mouse IgG antibodies (Invitrogen A31571, 1:100).

### Drug treatments

To inhibit ERK activation, U0126 (10 μM; MedChemExpress, HY-12031A) was added to M2 medium at different stages of oocyte maturation. Early U0126 treatment was performed during the first 6 h of maturation, before completion of spindle organization; Whereas late U0126 treatment was performed after spindle organization until PB1 extrusion, for approximately 6 h; Continuous U0126 treatment was performed from the GV stage until PB1 extrusion (Fig S4C). U0126 was prepared as a 50 mM stock solution in DMSO and diluted 1:5000 in M2 medium; control oocytes were treated with the same final concentration of DMSO. For mSIRK treatment, mSIRK (50 μM; MedChemExpress, HY-P5985) was diluted 1:1000 into M2 medium from a 50 mM stock solution.

### Imaging of oocyte

All high-resolution images of live oocytes were acquired using an Andor Dragonfly spinning-disk confocal microscope equipped with a 40× or 63×/1.44 NA oil-immersion objective, a scientific complementary metal-oxide semiconductor (sCMOS) camera (Andor Zyla 4.2), and an Andor Integrated Laser Engine (ILE) system with 405-, 488-, 568-, and 647-nm laser lines and a spinning-disk confocal scan head (Andor Dragonfly 500). The 405-nm laser was used for Hoechst, the 488-nm laser for mG, ZO-1-EGFP, and EGFP, the 568-nm laser for mT, and the 647-nm laser for Alexa Fluor 647 secondary antibody of immunofluorescent staining. During imaging, live oocytes were cultured in a mini-incubator mounted on the microscope stage. For transillumination imaging of live oocytes, images were acquired every 1 or 10 min. For *Oct4-CreER^T2^;mTmG* oocytes, images were acquired every 1 h; ZO-1-EGFP and EGFP-injected oocytes were imaged at interval 10 min. To track spindle movement trajectories while minimizing phototoxicity from the 405-nm laser, chromosome-labeled oocytes were imaged every 1 h using a 20× objective. All images were acquired using Fusion 2.1 software (https://andor.oxinst.com/products/dragonfly#fusion).

### Fluorescence recovery after photobleaching (FRAP)

In vitro FRAP experiments of ZO-1-EGFP were performed on a Leica STELLARIS 5 confocal microscope using a 488-nm excitation laser and 63× objective for both imaging and photobleaching. Images were acquired with Smart Gain set to 2.5%, Smart Intensity set to 0.5%, a zoom factor of 5, a scan speed of 400 Hz, and a frame size of 1024 × 1024 pixels. Selected ROIs were photobleached with 40% laser intensity, and the bleaching interval was set to the minimum value. Fluorescence recovery was recorded after bleaching using the same 488-nm imaging settings. Recovery curves were normalized to the fluorescence intensity of each ROI before photobleaching.

### Purification of proteins

Two recombinant mouse ZO-1 proteins were generated in this study: full-length ZO-1 for mass spectrometric analysis and full-length ZO-1 fused to C-terminal mEGFP and Flag tags (ZO-1-EGFP) for microinjection and LLPS assays.

Recombinant ZO-1 proteins were expressed in insect cells using the baculovirus expression system. *Spodoptera frugiperda* Sf9 cells infected with the corresponding recombinant baculovirus were cultured at 27°C in IB905 insect cell culture medium. At 60 h post-infection, cells were harvested, washed, resuspended in harvest buffer containing 25 mM HEPES, pH 7.4, 150 mM NaCl, and 1× protease inhibitor cocktail, and frozen in liquid nitrogen. All subsequent purification steps were performed at 4°C. Cells were lysed using a Dounce homogenizer, and the lysates were clarified by centrifugation at 50,000 × g for 35 min. The target protein was purified by immunoaffinity chromatography using anti-Flag M2 affinity resin. The resin was washed with 10 column volumes of wash buffer containing 25 mM HEPES, pH 7.4, and 500 mM NaCl, the protein was eluted with 0.4 mg/ml Flag peptide.

For purification of phosphorylated ERK2 (p-ERK2), Escherichia coli cells were co-transformed with plasmids encoding Strep-tagged ERK2 and His-tagged MEK1. Transformed cells were cultured at 37°C with shaking at 220 rpm until the OD_600_ reached 1.2, after which protein expression was induced with 0.5 mM IPTG. Cultures were then incubated at 18°C with shaking at 220 rpm for 16 h. Cells were harvested, washed, and resuspended in lysis buffer containing 25 mM Tris, pH 8.0, 150 mM NaCl, and 1 mM PMSF. Cells were lysed using a Dounce homogenizer, and lysates were clarified by centrifugation at 50,000 × g for 35 min. Strep-tagged p-ERK2 was purified by affinity chromatography using Strep-Tactin resin. After 10 column washes with wash buffer containing 25 mM Tris, pH 8.0, and 150 mM NaCl, the protein was eluted with 2.5 mM desthiobiotin.

### Phosphorylation assay of the ZO-1 and MS analysis

Purified ZO-1 protein (1.51 mg/ml) was subjected to in vitro phosphorylation using active p-ERK kinase. The reaction was performed by incubating ZO-1 with p-ERK (1.81 mg/ml) in buffer containing 25 mM Tris-HCl, pH 7.4, 1 mM DTT, 10 mM MgCl₂, 300 μM ATP, and 500 mM KCl at 30°C for 1 h. A parallel reaction without ATP served as a control. ZO-1 phosphorylation sites were then identified by mass spectrometry.

### MS analysis

Protein digestion was performed using FASP method with modifications (Nature Methods 6, 359 - 362 (2009). Briefly, 100 μg protein was dissolved with 50 mM ABC (NH_4_HCO_3_), reduced with DTT (dithiothreitol) at 56 ℃ for 45 min, and alkylated with IAM (iodoacetamide) at room temperature for 30 min in the dark. The solution was transferred into a 10K ultrafiltration tube (Vivacon 500, Satrorius), spinned at 14000 g for 20 min. 50 mM ABC solution was used to wash the protein for 3 times. 2 μg trypsin in 50 μL 50 mM ABC was added and incubated at 37 ℃ overnight. The ultrafiltration tube was spinned at 14000 g for 20 min with a new collection tube to collect digested peptides. ABC solution was added into the ultrafiltration tube to wash the digested peptide into the collection tube. The collected solution was diluted with 0.1% FA for nanoLC-MS analysis.

NanoLC separation was achieved with a Waters (Milford, MA, USA) nanoAcquityHPLC. The trap column was Thermo Acclaim PepMap 100 (75 μm × 2 mm, C18, 3 μm). The analytical column was homemade with 100 μm I.D. fused silica capillary (Polymicro) filled with 20 cm of C18 stationary phase (Phenomenex, Aqua 3 μm C18 125A). A gradient elution program was used, with mobile phase increases linearly from 1% B to 35% B in 65 min. Mobile phase A: 0.1% FA in water, B: 0.1% FA in acetonitrile.

Nanospray ESI-MS was performed on a ThermoOrbitrap Fusion Lumos high resolution mass spectrometer (Thermo Scientific, Waltham, MA, USA). With 120000 MS scan resolution and 15000 MSMS scan resolution. The full MS AGC target was 4E5 with an injection time of 50 ms and time between master scan is 3 seconds.

Raw data from the mass spectrometer were preprocessed with Mascot Distiller 2.7 for peak picking. The resulted peak lists were searched against Uniprot_Mouse (45,182 sequences; 20,089,089 residues) database using Mascot 2.7 search engine. The search parameters are: Fixed modifications: Carbamidomethyl (C), Variable modifications: Oxidation (M) and Phosphorylation (S, T, Y). Enzyme: Trypsin, Maximum missed cleavages: 2, MS mass tolerance: 10 ppm, MSMS mass tolerance: 0.02 Da. Scaffold PTM was used to evaluated phosphorylation sites of the Mascot search results using Ascorealgorithm.

### In vitro phase-separation of ZO-1-EGFP

Purified ZO-1-EGFP (5.21 mg/mL) was incubated with active p-ERK2 (3.60 mg/mL) in phosphorylation buffer containing 200 μM ATP at 30 °C for 1 h. Parallel reactions lacking ATP and/or p-ERK2 were included as controls. The reactions were subsequently diluted into phase-separation buffer containing 25 mM HEPES (pH 7.4), 150 mM KCl, and 2% PEG8000, yielding final ZO-1-EGFP and p-ERK2 concentrations of approximately 9.77 and 2.14 μM, respectively. Droplet formation was examined by fluorescence microscopy using 488-nm excitation under identical imaging conditions.

### Microinjection of mouse oocytes

Purified ZO-1-EGFP protein (4 mg/ml) was microinjected into oocytes, and purified EGFP protein (0.996 mg/ml) was used as a control. Denuded oocytes were microinjected in M2 medium supplemented with 2.5 μM milrinone. Microinjection needles were pulled from borosilicate glass capillaries (item no. BF100-78-10). Approximately 10 pL protein was injected into each oocyte using a FemtoJet 4× electric microinjector (Eppendorf), followed by immediate imaging with an Andor Dragonfly spinning-disk confocal microscope using the parameters described above.

### Oocyte modeling and analysis

Three-dimensional reconstruction and modeling of oocytes were performed using Imaris software. For membrane-labeled oocytes from *Oct4-CreER^T2^;mTmG* ovaries, whole-oocyte surface models were generated by detecting membrane fluorescence intensity, allowing quantification of surface area, volume, and displacement over time. Regional membrane models were generated from manually selected ROIs.

ZO-1-EGFP puncta were detected based on fluorescence intensity within selected ROIs, with identical detection parameters applied to different ROIs from the same oocyte. Because ZO-1-EGFP puncta were distributed beneath the oocyte membrane, membrane surfaces in ROIs containing puncta were reconstructed through iterative machine-learning-based segmentation and manual correction guided by ZO-1-EGFP fluorescence signals.

For spindle-size analysis, spindle structures in control and U0126-treated oocytes were reconstructed directly from antibody fluorescence intensity in immunofluorescence images using intensity-based modeling.

Data generated from all modeling analyses were exported, analyzed, and plotted using Origin software.

### Western blot

Purified protein samples were separated using 12% SDS-PAGE gels (ACE Biotechnology, ET10012Gel) and subsequently transferred to PVDF membranes (IPVH00010; Millipore). Membranes were blocked in TBST containing 5% BSA for 1 h at room temperature, and then incubated overnight at 4°C with the indicated primary antibodies: mouse anti-Flag antibody for detection of ZO-1 and ZO-1-EGFP (Mouse, 1:5000; CWBIO, CW0287M) and anti-Strep antibody for ERK2 detection (Rabbit, 1:1000; Abcam, ab27591). Membranes were washed three times with TBST and incubated with HRP-conjugated secondary antibodies (1:5000; Beyotime) for 1 h at room temperature.

### Image analysis and quantification

Following image acquisition, movies and single time-point images were processed in ImageJ (http://rsbweb.nih.gov/ij/) for z-stack projection and channel merging. In Supplementary Fig. 2, the 647-nm channel was inverted to black and white in ImageJ to enhance visualization of cortical actin structures. Rotating 3D supplementary movies illustrating oocyte membrane movement and selected models were processed using Imaris software (https://imaris.oxinst.com/) (Movies S1, S2, S3, S4 and S5).

### Comb-like structures quantification and analysis

Comb-like structures (Oo-comb) were identified by layer-by-layer examination of z-slices at each time point. Unlike Oo-Mvi, Oo-comb were defined as straight structures lacking spherical vesicles and not invading the zona pellucida after PES formation. Oo-comb and Oo-Mvi were manually counted in selected ROIs at each time point using ImageJ. For normalization, the number of structures at the initial time point was set to 100%, and counts at subsequent time points were normalized to the initial value.

### Spindle quantification and analysis

To assess the relationship between PES location and spindle migration direction, the spindle migration angle α was defined as the angle between two lines: line m, which connects the edge of the PES with the center of the oocyte, and line n, which passes through the two spindle poles (Fig.2D). When the line n toward in PES region, the angle α set to 0. In addition to the initial spindle migration direction, a second determinant of spindle trajectory is spindle rotation after cortical arrival. Spindles were classified as moving toward or deviating from the PES based on the rotation angle α (Fig. 2F).In the U0126 treatment assay, the minimum distance between the chromosome cluster and the cortex was used as an indicator of spindle migration toward the cortex.

### PES quantification and analysis

The areas of the PES and PB1 were quantified using FIJI (ImageJ) from transillumination images of oocytes. For each oocyte, the relevant z-slice showing the maximal cross-sectional area was selected for analysis. The maximum PES distance was defined as the greatest linear distance between the oocyte plasma membrane and the zona pellucida within the PES region. This distance was measured using the straight-line tool in FIJI on the same representative slice used for area quantification.

All experiments were repeated at least three times, Data are presented as the mean ± standard deviation (SD) of each experiment. Statistical analyses and graph generation were performed using Prism v10.6.1 (GraphPad Software, La Jolla, CA, USA) and Origin. Comparisons between two independent groups were analyzed using a two-tailed unpaired Student’s t-test (Figures 1B, 1H, 3D, 4C, 4E, 4F, and 5B, 5C; Supplementary Figures 2C and 4B). Paired comparisons were analyzed using a paired t-test (Figure 3G and Supplementary Figure 3C). Comparisons among multiple groups were performed using one-way ANOVA followed by Tukey’s multiple-comparisons test (Figures 1C, 1I, 3E, 3K, 4J, 5E, 5F, and 5I; Supplementary Figure 4C). Statistical significance was set at p < 0.05. Significance levels are indicated as follows: *p < 0.05, **p < 0.01, ***p < 0.001; ns, not significant, p ≥ 0.05.

---

**Supplementary Fig. 1.**
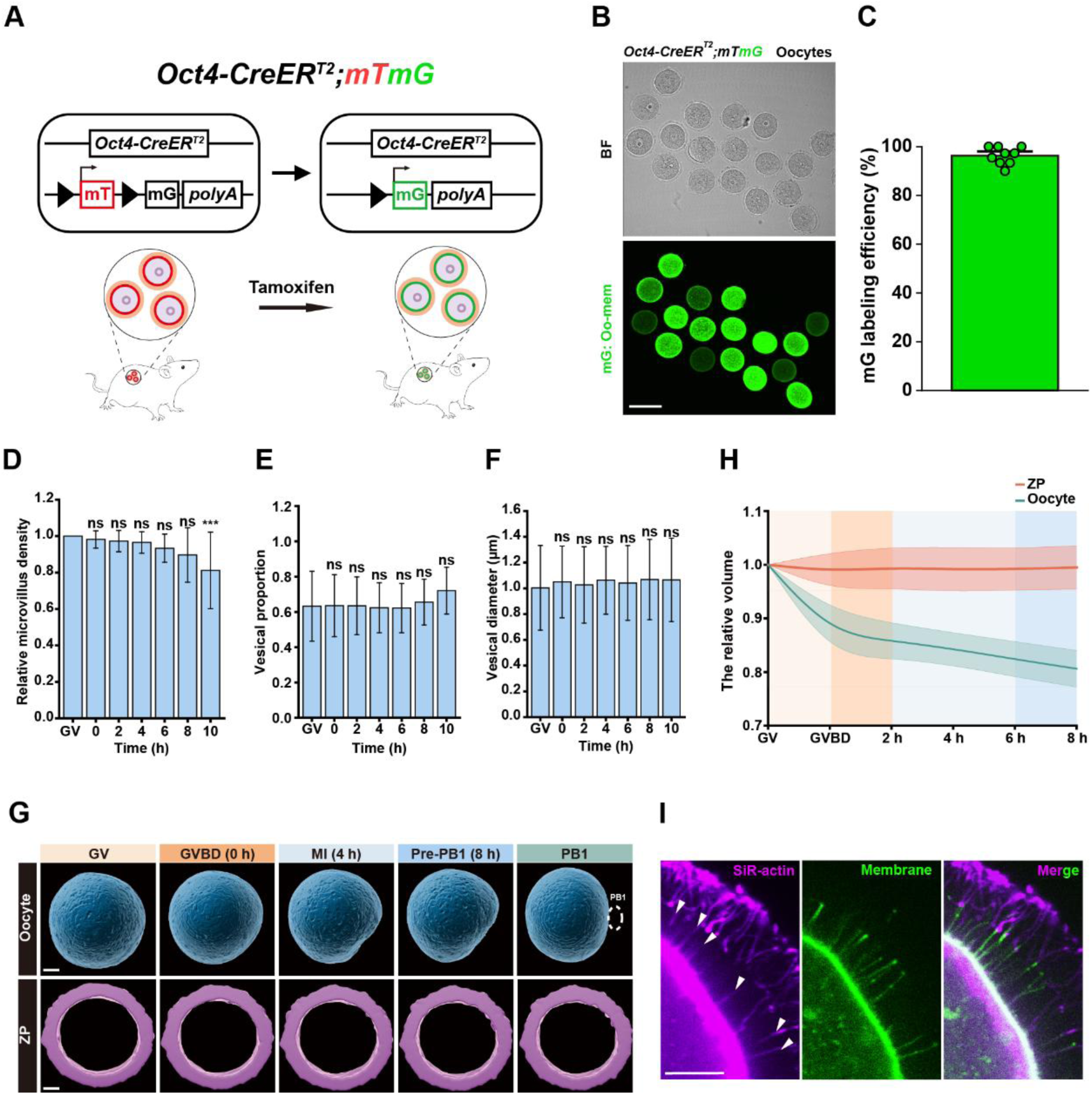
Genetic membrane labeling and three-dimensional analysis of oocyte and zona pellucida volume during maturation. A. Schematic of the strategy used to label the oocyte membrane in *Oct4-CreER^T2^;mTmG* mice. Membrane-localized fluorescent protein (mT) switches to green-fluorescent protein (mG) following oocyte-specific expression of *CreER^T2^* recombinase induced by tamoxifen administration. B. Representative bright field and fluorescence images of oocytes isolated from *Oct4-CreER^T2^;mTmG* ovaries show uniform green fluorescence labeling of the oocyte membrane. C. Quantification of membrane-GFP labeling efficiency, expressed as the percentage of GFP-positive oocytes obtained from each mouse (n = 9 mice). D. Quantification of relative Oo-Mvi density during oocyte maturation (n = 18 oocytes). Density did not differ significantly from GV at 0–8 h but decreased significantly at 10 h. E. Quantification of the proportion of Oo-Mvi associated with vesicles during oocyte maturation (n = 16 oocytes). No significant overall difference was detected across maturation stages. F. Quantification of the diameter of vesicles associated with Oo-Mvi during oocyte maturation. A total of 299 vesicles from 15 oocytes were analyzed. Sample sizes were 43, 45, 46, 49, 43, 39 and 34 vesicles at GV, 0, 2, 4, 6, 8 and 10 h, respectively. No significant overall difference in vesicle diameter was detected. G. Representative three-dimensional time-lapse models of the oocyte cell body and ZP during meiotic maturation, showing a progressive reduction in oocyte volume. The extruded PB1 is outlined by the white dashed circle. H. Quantification of oocyte and ZP volumes during maturation. Oocyte volume decreases significantly after GVBD, whereas ZP volume remains largely unchanged (n = 20 oocytes). GVBD was defined as 0 h. Relative volume was normalized to the volume analyzed at GV. I. Representative images of a membrane-GFP-labelled oocyte stained with SiR-actin at 4 h after GVBD. White arrowheads indicate F-actin-containing Oo-Combs. Scale bars, 100 μm (B) and 10 μm (F and H). Data in C, D, E, F and H are shown as mean ± s.d. ns, not significant. ***P < 0.001. Statistical significance was assessed using one-way repeated-measures ANOVA in D, E and F.

**Supplementary Fig. 2.**
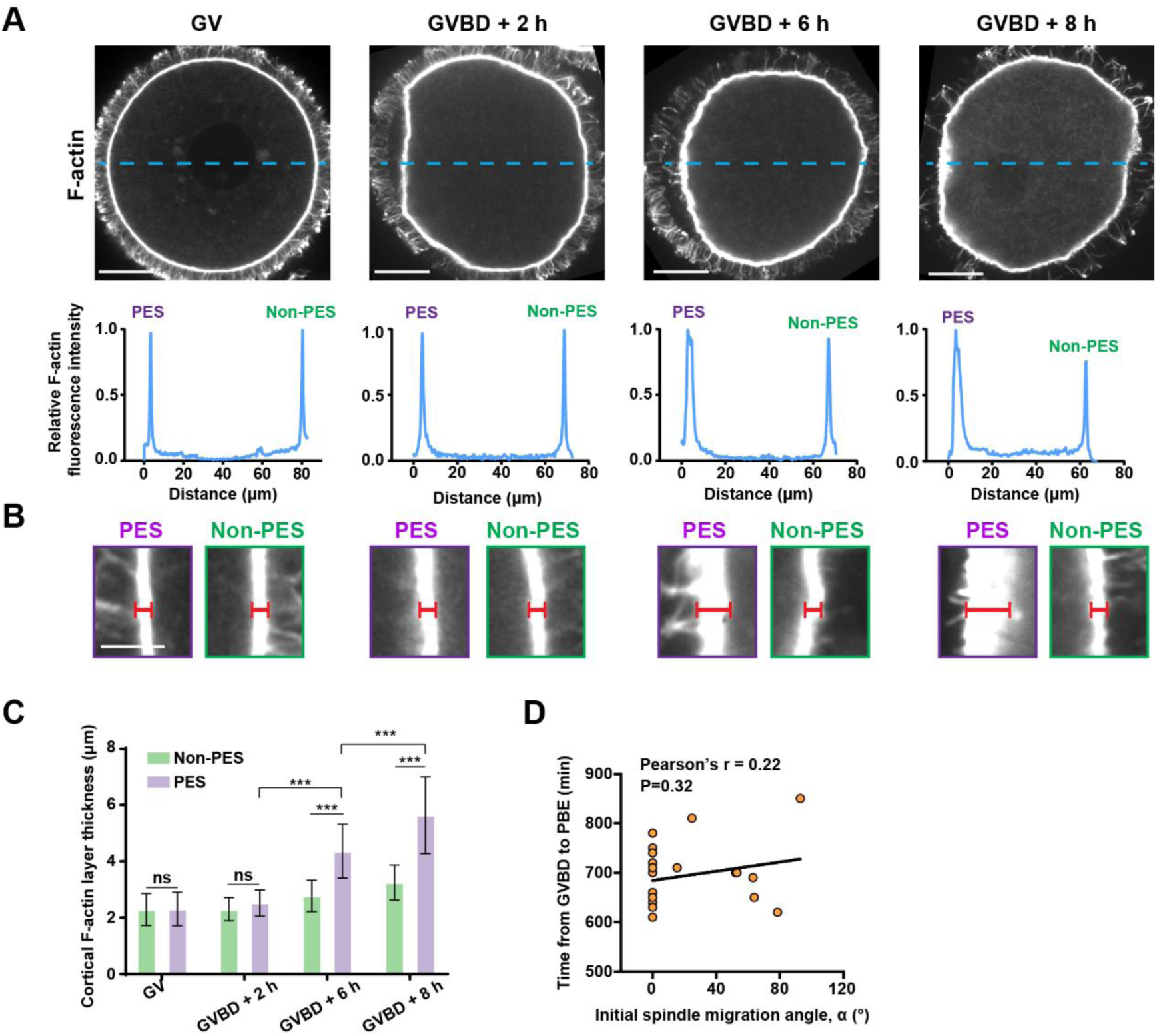
PES formation precedes localized cortical-actin thickening. A. Representative images of phalloidin-stained oocytes at the indicated stages of maturation (top) and corresponding cortical F-actin fluorescence-intensity profiles from the PES-associated and non-PES regions (bottom). Blue dashed lines indicate the line-scan paths. B. Magnified views of the PES-associated and non-PES cortical regions shown in A. Red lines indicate the cortical-actin thickness measurements. C. Quantification of cortical-actin thickness in the PES-associated and non-PES regions during oocyte maturation. Cortical-actin thickness did not increase detectably during the initial formation of the PES but became significantly elevated in the PES region at approximately 6 h after GVBD. Cortical thickness in the non-PES region remained comparatively stable throughout maturation (n = 20, 21, 13 and 11 oocytes at each time, respectively). D. Correlation between the initial spindle-migration angle α and the time of PB1 extrusion. Initial spindle direction was not significantly associated with the overall timing of PB1 extrusion (n = 19 oocytes). Each point represents one oocyte. Scale bars, 20 μm in A and 5 μm in B. Data in C are shown as mean ± s.d.; ns, not significant; ***P < 0.001. Comparisons between regions at each time point used two-tailed paired Student’s t-tests, and comparisons across time within each region used one-way ANOVA. The correlation in D was assessed using two-tailed Pearson’s correlation analysis.

**Supplementary Fig. 3.**
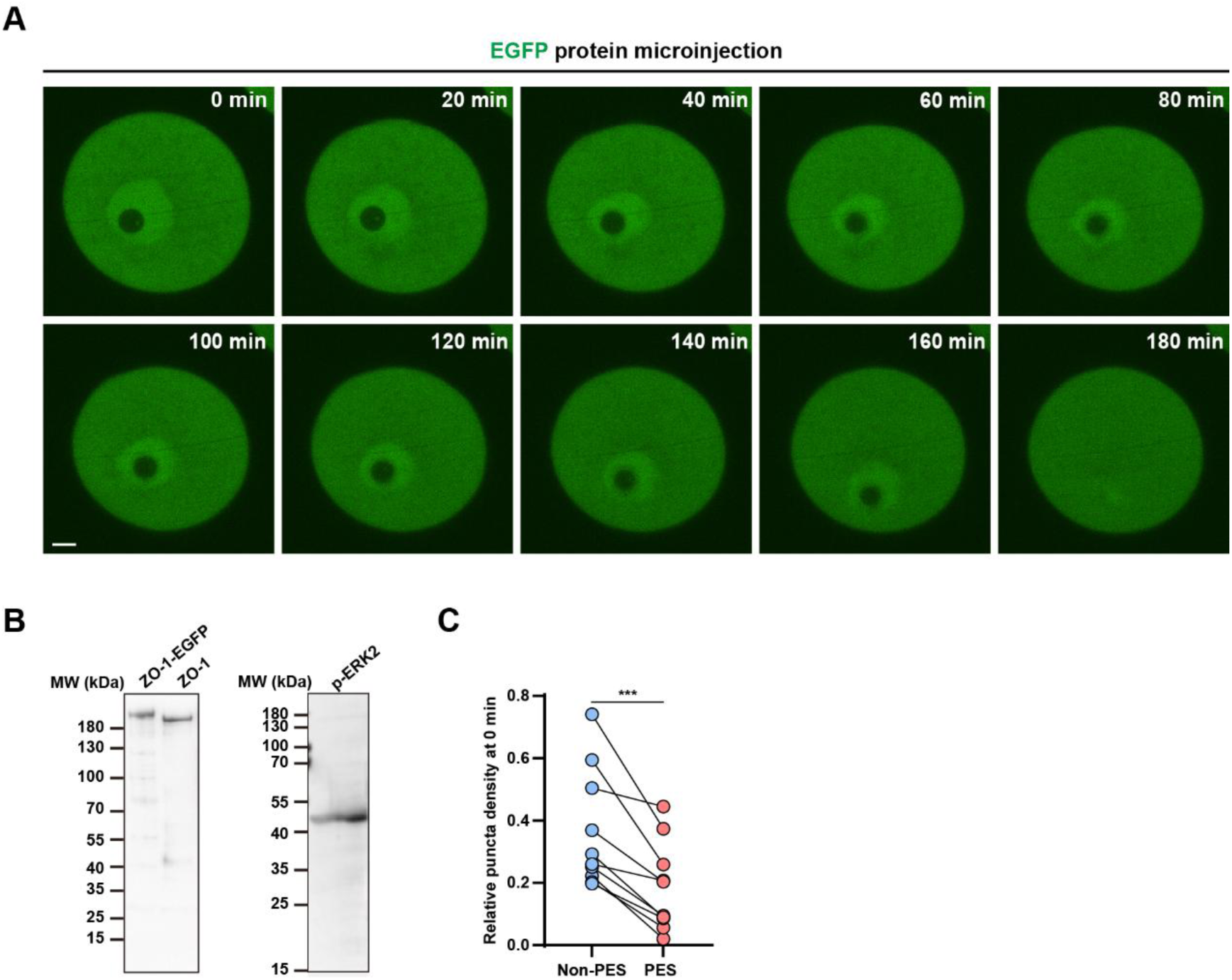
Validation of recombinant ZO-1 and active ERK2 proteins. A. Representative images of a GV-stage oocyte microinjected with mEGFP alone, showing uniform cytoplasmic fluorescence without membrane-associated puncta. The time of capturing first picture of oocyte was defined as 0 min. B. Immunoblot analysis of purified ZO-1-EGFP, ZO-1 and active p-ERK2 proteins used for microinjection and in vitro biochemical assays. C. Paired quantification of ZO-1-EGFP puncta area density at the beginning of time-lapse imaging. The membrane region that subsequently formed the PES (PES) exhibited lower puncta density than the non-PES region within the same oocyte (n = 10 oocytes). Lines connect PES and non-PES measurements from the same oocyte. Scale bar is 10 μm (A). ***P < 0.001 by two-tailed paired Student’s t-test in C.

**Supplementary Fig. 4.**
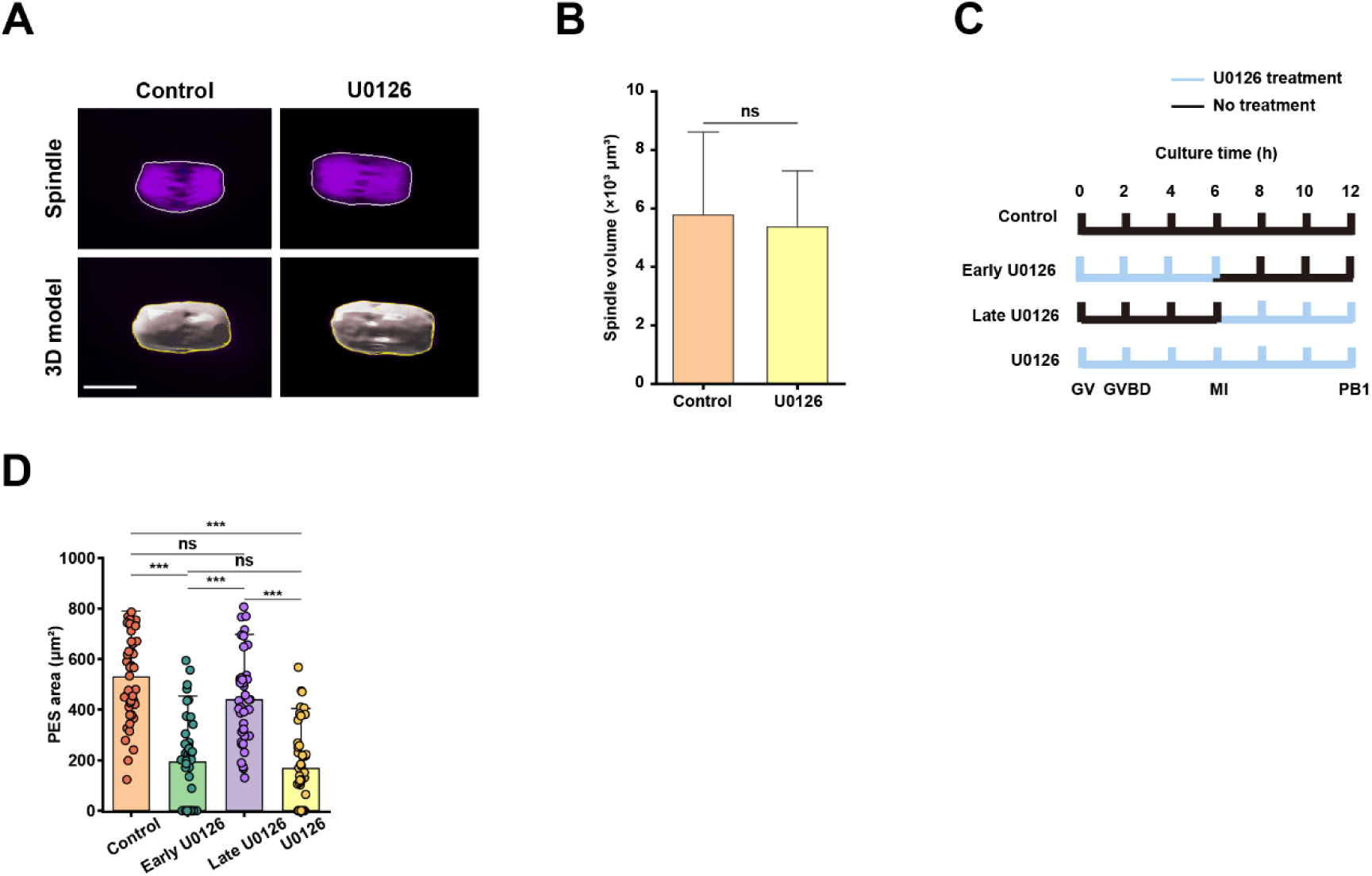
Effects of U0126 treatment on spindle organization and PES formation. A. Representative α-tubulin immunofluorescence images and corresponding three-dimensional spindle models from control and U0126-treated oocytes, showing no overt difference in spindle organization. B. Quantification of spindle volume based on three-dimensional reconstruction, showing no significant difference between control and U0126-treated oocytes (n = 20 oocytes for each group). C. Schematic of the continuous, early and late U0126 treatment regimens. Blue lines: the periods of U0126 treatment; Black lines: the periods of culture in inhibitor-free medium. D. Quantification of PES area following the indicated U0126 treatment regimens. Continuous and early U0126 treatment reduce PES area, whereas late U0126 treatment has no significant effect compared with the untreated control (control, n = 43 oocytes; Early U0126, n = 42 oocytes; Late U0126, n = 44 oocytes; U0126, n = 45 oocytes). Scale bar, 20 μm (A). Data are shown as mean ± s.d.; ns, not significant; *P < 0.05, **P < 0.01, ***P < 0.001 by two-tailed unpaired Student’s t-test (B) and one-way ANOVA followed by Tukey’s multiple-comparisons test (C).

**Supplementary Fig. 5.**
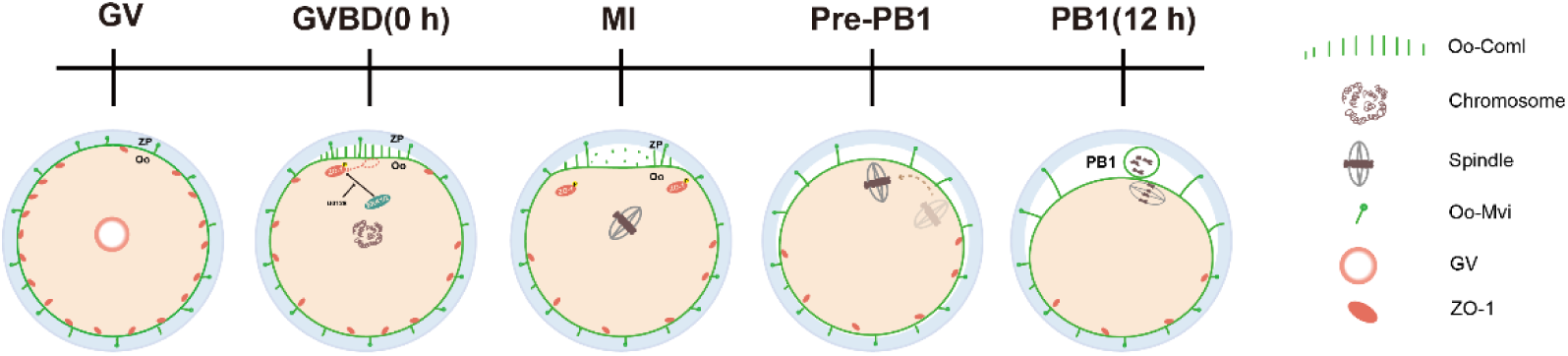
Schematic illustration of membrane pre-remodeling event and PES-associated preferentially location of spindle before PB1 extrusion. At the GV stage, ZO-1 puncta are heterogeneously distributed in oocytes, with fewer puncta in the prospective PES region. Around GVBD, ERK-dependent phosphorylation suppresses ZO-1 phase separation, rapidly weakening the oocyte-zona pellucida association in the PES region. The oocyte consequently detaches locally from the zona pellucida to form the PES. Following MI spindle assembly, migration occurs in two steps. The initial trajectory toward the cortex is random relative to the PES. After reaching the cortex, the spindle preferentially repositions toward the PES, where PB1 is subsequently extruded.

## Acknowledgements

The authors are grateful to Prof. Guanghui Yang for support with protein purification and related experiments; Rong Wang and Profs. Yan Guo and Jingrui Li for assistance with phase separation experiments; and Yan Li and Prof. Xuguang Du for support with microinjection experiments. We appreciate the assistance provided by the biological mass spectrometry laboratory from College of Biological Sciences at China Agricultural University on our mass spectrometry experiment. This study was supported by the National Key Research and Development Program of China to Y.Z., H.Z. and J.L. (2024YFC2706600); The National Natural Science Foundation of China to Y.Z. (82371664, 32541050). The Key Program of National Natural Science Foundation of China to H.Z. (82230051); Beijing Natural Science Foundation to H.Z. and Y.Z. (L2609048 and 5262014) and The 2115 Talent Development Program of China Agricultural University to Y.Z and H.Z. (1021-00109022 and 1021-15055007).

## Contributions

K.G., Y.Z., and H.Z. designed the research; K.G., X.H., Y.B., P.F., N.S., Z.W., and C.Y. performed the experiments; K.G., X.H., G.W., X.Y., J.L., K.C., G.Y., Y.Z., Z.L., and H.Z. analyzed the data; K.G. draw the graphic model. K.G., H.Z., and Y.Z. wrote the paper. All authors have seen and approved the final version.

## Compliance and ethics

The authors have declared that no conflict of interest exists.

